# The role of evolutionarily metastable oligomeric states in the optimization of catalytic activity

**DOI:** 10.1101/2022.09.13.507756

**Authors:** Nicholas J. East, Ben E. Clifton, Colin J. Jackson, Joe A. Kaczmarski

**Affiliations:** ARC Centre of Excellence in Synthetic Biology, Australian National University, Canberra, Australia; Research School of Biology, Australian National University. R.N Robertson Building, Building 46, Biology Place, Acton, Australian Capital Territory, Australia 2601; ARC Centre of Excellence for Innovations in Peptide and Protein Science, Research School of Chemistry, Australian National University. Building 137, Sullivans Creek Rd, Acton, Australian Capital Territory, Australia 2601; Protein Engineering and Evolution Unit, Okinawa Institute of Science and Technology, 1919-1 Tancha, Onna, Okinawa, Japan 904-0412

**Keywords:** Oligomerization, protein evolution, cyclohexadienyl dehydratase, enzyme evolution, size-exclusion chromatography, molecular dynamics simulations, trimerization

## Abstract

The emergence of oligomers is common during the evolution and diversification of protein families, yet the selective advantage of oligomerization is often cryptic or unclear. Oligomerization can involve the formation of isologous head-to-head interfaces (e.g., in symmetrical dimers) or heterologous head-to-tail interfaces (e.g., in cyclic complexes), the latter of which is less well studied and understood. In this work, we retrace the emergence of the trimeric form of cyclohexadienyl dehydratase from *Pseudomonas aeruginosa* (PaCDT) by introducing residues that form the PaCDT trimer-interface into AncCDT-5 (a monomeric reconstructed ancestor of PaCDT). We find that single interface mutations can switch the oligomeric state of the variants (implying evolutionarily metastable oligomeric states) and that trimerization corresponds with a reduction in the *K*_M_ value of the enzyme from a promiscuous level to the physiologically relevant range. We show that this can be rationalized at the structural and dynamic level by reduced sampling of a non-catalytic conformational substate, and that trimerization was likely followed by a C-terminal extension that further refined the conformational sampling and kinetic properties of the enzyme. This work provides insight into how neutral sampling of metastable oligomeric states along an evolutionary trajectory can facilitate the evolution and optimization of enzyme function.

**Importance & Impact Statement:** Understanding how and why structural complexity (including homo-oligomerization and sequence insertions) emerges during the evolution and diversification of natural enzymes is a key goal in the study and design of protein function. We show that cyclic homo-oligomeric states can emerge via a small number of substitutions, and that trimerization and a C-terminal extension contributed to the tuning of catalytic properties during the evolution of cyclohexadienyl dehydratase from *Pseudomonas aeruginosa*.

## Introduction

A large proportion of proteins self-associate to form homo-oligomers comprising two or more identical subunits.^1–4^ Among other things, homo-oligomerization has been shown to play an important role in conferring gain of function^5,6^, protection from degradation^7^, cooperative binding properties^8^, allosteric regulation of enzyme activity^9^, and enhanced thermostability of proteins^10,11^. On the other hand, Lynch and others have demonstrated that oligomers can also arise and become entrenched even when there is no apparent adaptive advantage associated with the initial formation of the complex.^12–14^ Considering the prevalence and biological importance of homo-oligomers, understanding the evolutionary factors that drive the formation of new homo-oligomeric species and the effect that oligomerization has on protein properties are key goals in the study and design of protein complexes.^15–19^

In order to probe the drivers, mechanisms and functional consequences of oligomerization, several studies have used evolution-based approaches to investigate how the initial emergence and/or diversification of oligomeric states affected protein properties during the natural or laboratory-based evolution of proteins.^12^ For example, ancestral sequence reconstruction (ASR) revealed that two historical substitutions conferred a switch between a dimeric and tetrameric form during the evolution of vertebrate haemoglobin, and that this shift in oligomeric state gave rise to beneficial cooperative binding of oxygen.^8^ Similarly, ASR revealed the role of quaternary structure plasticity during the evolution and diversification of Ribulose-1,5-bisphosphate carboxylase-oxygenase (RuBisCO) function.^6^ Shifts in oligomeric states have also been linked to enhanced thermostability during the laboratory-directed evolution of an αE7 carboxylesterase^10^ and the tailoring of activity and structural stability amongst bacterial methionine S-adenosyltransferase (MAT) homologs.^20^ On the other hand, a dimerization event that occurred during the evolution of steroid hormone receptors provided no immediate selective advantage.^14^ While evolution-based studies such as these have provided valuable insight into the mechanisms and functional importance of historical dimerization events (leading to proteins with C2 or D2 symmetry, for example), less is known about the emergence of cyclic complexes such as homotrimers.

Cyclohexadienyl dehydratase from *Pseudomonas aeruginosa* (PaCDT) is a homotrimeric enzyme that evolved from a monomeric, non-catalytic ancestral solute-binding protein (**Figure 1A**).^21,22^ In our previous work, we observed a ∼60-fold increase in catalytic efficiency between the monomeric AncCDT-5 (the most recent ancestor of PaCDT that we characterized) and modern PaCDT, despite these two proteins sharing identical active site residues.^21,23^ Interestingly, while the subunits of PaCDT and its ancestors share the same bilobed periplasmic binding protein-like II fold (comprising two subdomains linked by a flexible hinge region), we found that the monomeric ancestors of PaCDT, including AncCDT-5, mostly sampled open or wide-open states (in which their two sub-domains were far apart), while trimeric PaCDT predominantly samples a more compact, and catalytically-relevant, closed state (in which the two domains are closer together and the active site is pre-organized for catalysis). We speculated that the shift in open-closed sampling and the associated increase in dehydratase activity between AncCDT-5 and PaCDT may have been, at least in part, due to oligomerization of the protein restricting rigid-body motions of the individual chains.

**Figure 1.**
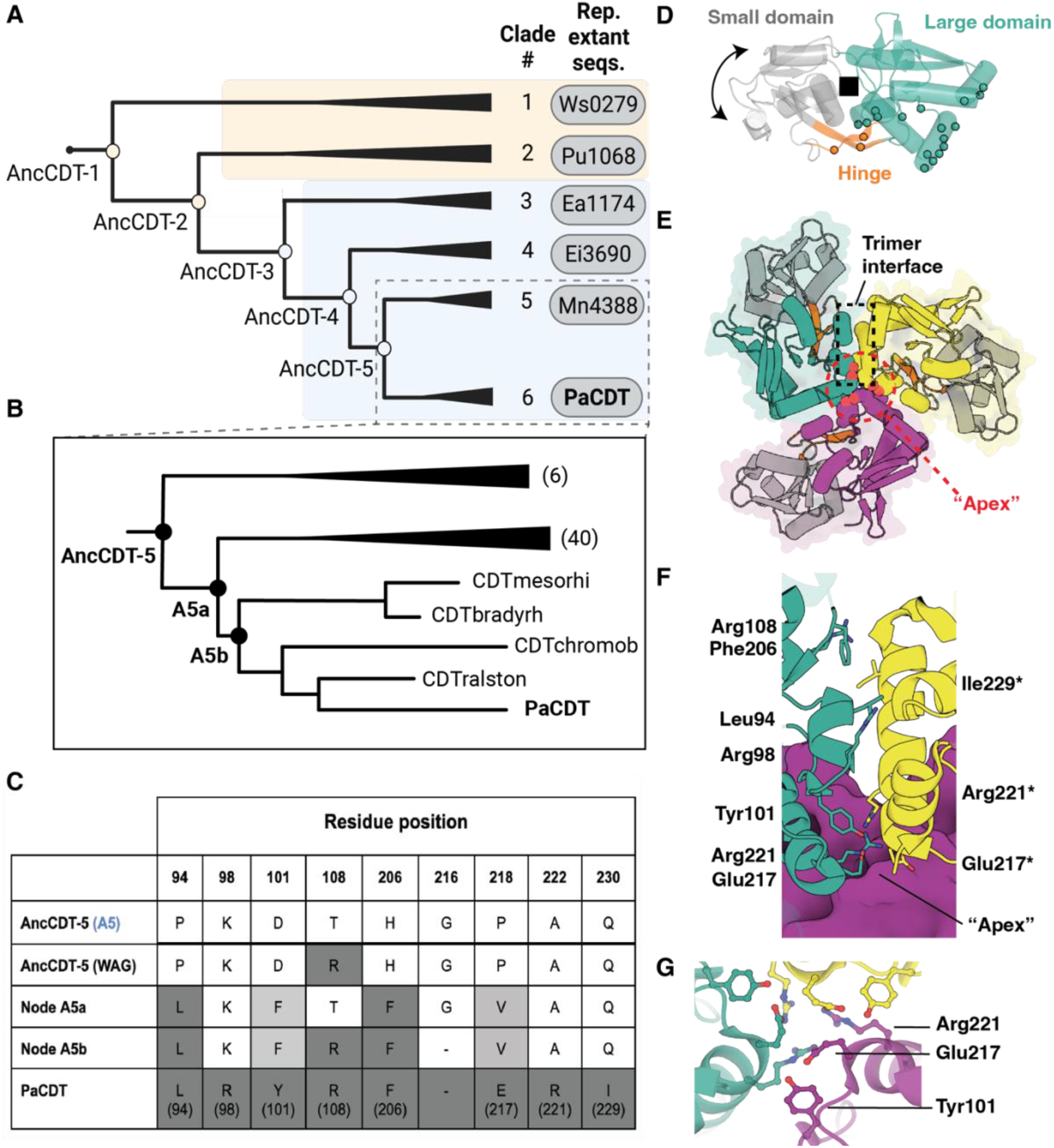
Design of the trimer-interface variants of AncCDT-5. **(A)** A simplified version of the phylogenetic tree used in our previous work^21^ to explore the evolution of PaCDT. Key ancestral nodes (AncCDT-1 to AncCDT-5) are labelled. Representative extant proteins are shown next to the clades in which they are found: Ws0279 (UniProt: Q7MAG0), Pu1068 (UniProt: Q4FLR5), Ea1174 (UniProt: K0ABP5), Ei3690 (UniProt: C5B978), Mn4388 (UniProt: B8IBK5). **(B)** A zoom-in of the phylogenetic tree shown in (A), highlighting the extant CDT homologs and internal ancestral nodes between AncCDT-5 and PaCDT that were used to guide the design of the AncCDT-5 trimer-interface mutants. The number of sequences in each collapsed clade is shown in parentheses. Extant sequences: CDTmesorhi (UniProt: I2F416), CDTbradyrh (UniProt: A4YUK0), CDTchromob (UniProt: Q7P1I9), CDTralston (UniProt: Q8XWG2). **(C)** Table showing residues of AncCDT-5, PaCDT and internal ancestral nodes at positions at the PaCDT trimer interface that differ between AncCDT-5 and PaCDT. The residues at the equivalent positions in an alternate reconstruction of AncCDT-5 (AncCDT-5 (WAG)^21^) are also shown. The residue numbering is based on that used for the crystal structure of AncCDT-5 (PDB 6WUP); PaCDT residue numbers (according to PDB 5HPQ) are shown in parentheses. **(D-G)** The crystal structure of PaCDT (PDB 5HPQ) and trimer interface residues. **(D)** A single chain from the crystal structure of PaCDT showing the small (grey) and large (teal) subdomains, and the flexible hinge region (orange). The positions of residues that form the trimer interface are indicated by spheres. The active site region is shown as a black square. **(E)** The trimeric form of PaCDT, showing the three chains interacting mainly via the large domain of each chain. **(f)** Key trimer interface residues that differ between AncCDT-5 and PaCDT shown as sticks (numbered according to PDB 5HPQ). **(G)** Key residue positions at the apex of the trimer in PaCDT.

In this work, we used crystal structures of PaCDT and the sequences of modern descendants of AncCDT-5 to guide the design of a series of variants of AncCDT-5 that differed only at positions that form the trimer interface in PaCDT. By doing so, we (i) discovered two distinct mutational pathways linking the ancestral monomeric AncCDT-5 with higher oligomeric states, (ii) identified variants that differ by only one or two substitutions at the trimer interface but that have distinct oligomeric states, (iii) show that oligomerization results in a decrease in the *K*_M_ of the enzyme, bringing it close to physiologically relevant concentrations of the substrate and (iv) show that this is consistent with a shift in the conformational landscape towards more catalytically-relevant states. Finally, we demonstrate that the subsequent gain of a C-terminal extension further refines the conformational sampling and activity of the enzyme.

## Results

### Designing trimer-interface variants of AncCDT-5

In our previous work, we hypothesized that the trimeric nature of PaCDT may have contributed to the >50-fold greater dehydratase catalytic efficiency, compared with its monomeric ancestors, including AncCDT-5.^**21**,**23**^ The PaCDT trimer is formed through interactions between the larger of the two sub-domains in each of the three chains, as well as some residues in the flexible “hinge” region of each chain (**Figure 1D-G**). The three chains of the trimer are arranged around the point of three-fold symmetry (i.e., C3 symmetry) somewhat like a three-pronged pinwheel, with the larger sub-domains of the three chains coming together near the “apex” of the trimer (**Figure 1E**). Since the smaller sub-domains are not part of the trimer interface, this arrangement allows for some rigid-body open-closed motions to occur in each of the chains. However, we reasoned that the packing of the chains in the trimer might sterically limit or restrict the sampling of wide-open conformations and lead to enhanced activity due to increased sampling of the catalytically-competent closed state.^**23**^

To determine if trimerization of the protein alone could have contributed to the increase in catalytic efficiency that we previously observed between AncCDT-5 and PaCDT (which differ at 88 residue positions),^21,23^ we generated variants of AncCDT-5 that had different oligomeric states but differed only by substitutions at the trimer-forming interface. Sequence and structural alignments of PaCDT (PDB 5HPQ) and AncCDT-5 (PDB 6WUP) revealed eight key positions spread across the trimer-forming interface that differed between AncCDT-5 and PaCDT (**Figure 1C,F**). The substitution of residues at the apex positions of the PaCDT trimer (including PaCDT residues Tyr101, Glu217 and Arg221, hereafter termed “YER”) were of particular interest (**Figure 1G**); we expected that these residues would likely be important for determining whether three chains could pack together to form the trimeric structure. Interestingly, the sequences of close homologs of PaCDT (i.e., the extant sequences found in the clades descendent from AncCDT-5) showed variation at these three positions; for example, CDT homologs from *Bradyrhizobium* sp. ORS 278 (UniProt A4YUK0) and *Ralstonia solanacearum* CMR15 (UniProt D8N7P8) both have residues Phe101 and Val218 at the apex rather than the residues found in PaCDT (Tyr101, Glu217) or AncCDT-5 (Asp101, Pro218) (**Supplementary Table I**).

To identify substitutions that would switch the monomeric AncCDT-5 to a trimer, we designed a series of variants that introduced the PaCDT trimer interface-forming residues into AncCDT-5 in a stepwise manner, with most consecutive variants differing by only one or two mutations at the trimer interface. The order in which the substitutions were introduced was guided by the sequences of the present-day descendants of AncCDT-5 and by inferring the most likely ancestral sequences at internal nodes in the phylogenetic tree between AncCDT-5 and PaCDT (“A5a” and “A5b”, **Figure 1B,C**). For example, since all extant descendants of AncCDT-5 had Leu94 and Phe206, the most parsimonious explanation is that mutations P94L and H206F were introduced earlier in the evolutionary sequence.

Considering that some of the descendants of AncCDT-5 had Val217 or Phe101 at the apex positions and that reconstruction of ancestral nodes suggested that these residues may have been present in intermediate ancestral states (**Figure 1C**), we also created variants that introduced these substitutions into AncCDT-5. Overall, the series of trimer-interface variants (**Figure 2A**) culminated in an AncCDT-5 variant that shared all trimer interface residues with PaCDT (A5+CDT_interface_). We also generated a variant of AncCDT-5 that contained all eight trimer-interface residues identified in PaCDT and had Gly216 removed (A5+CDT_interface_ΔG216); PaCDT has a deletion at this position, which is located near the apex of the trimer. Plasmids (pET28a) containing sequences encoding N-terminal His-tagged PaCDT, AncCDT-5 and the trimer-interface AncCDT-5 variants were produced. All proteins expressed in their soluble, folded state in *Escherichia coli*.

**Figure 2.**
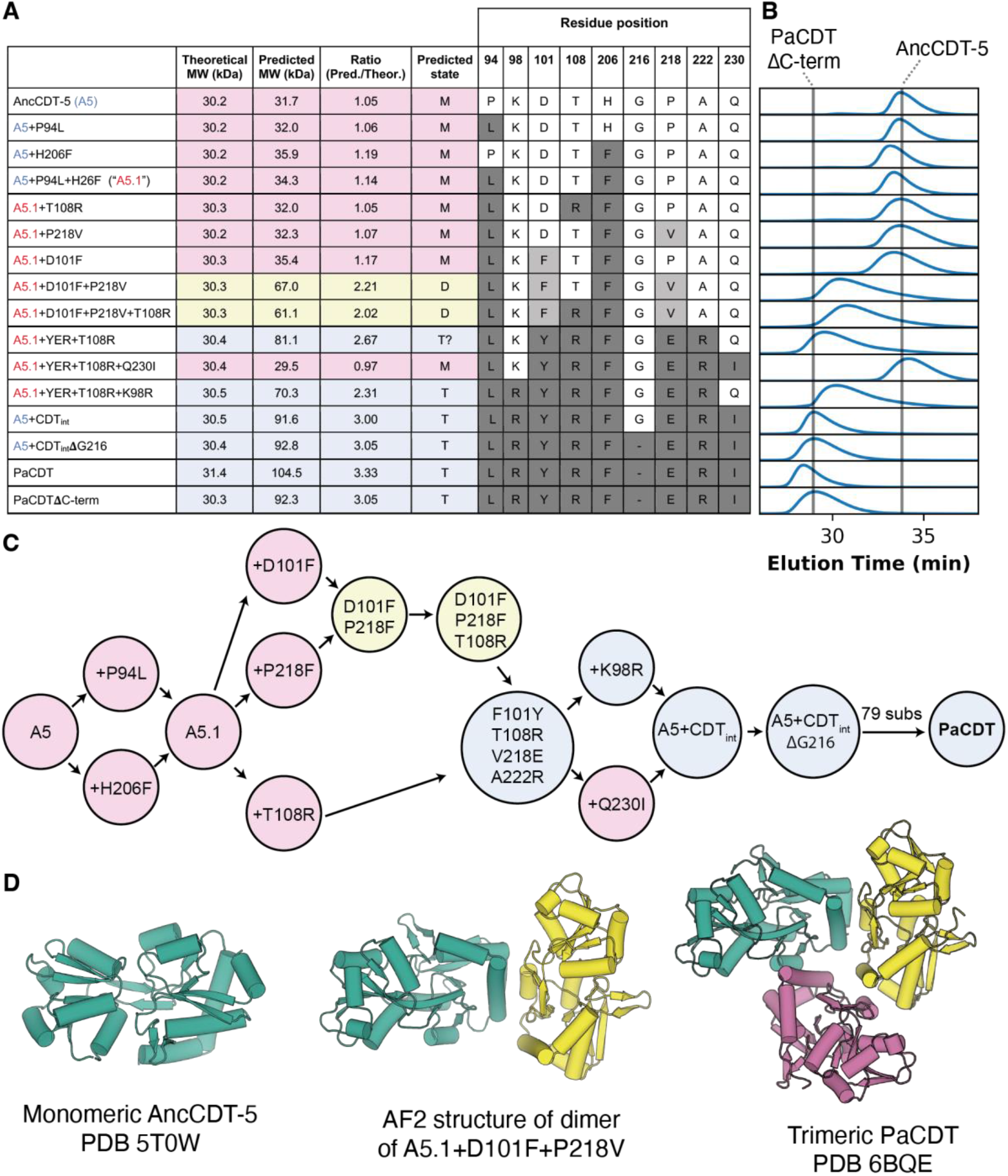
Oligomeric states of trimer-interface variants of AncCDT-5. **(A)** Table showing residues in trimer-interface variants, predicted molecular weight (MW) and oligomeric state determined by size-exclusion chromatography (M=monomer, D=dimer, T=trimer). Proteins were injected onto the SEC column at 8 mg/mL. Associated SEC-MALLS data is provided in (**Supplementary Figure 1**). Residues found in PaCDT are shaded dark grey. Residues not found in either AncCDT-5 or PaCDT are shaded light-grey. **(B)** Normalized refractive index chromatograms showing elution peaks of trimer-interface variants. Vertical lines aligned with elution peaks of -AncCDT-5 and PaCDTΔC are shown for reference. **(C)** Schematic showing mutational pathways linking AncCDT-5 and PaCDT via the interface variants in this study (colored by predicted oligomeric state). **(D)** From left to right, structures of the monomeric (AncCDT-5, PDB 5T0W), dimeric (Alpha-Fold2 model of A5.1+D101F+P218V) and trimeric (PaCDT, PDB 6BQE) forms of the related proteins.

### Oligomeric structures of AncCDT-5 trimer-interface variants

The oligomeric states of purified AncCDT-5, PaCDT and several key AncCDT-5 trimer-interface variants were determined by analytical size exclusion chromatography (SEC) and/or SEC coupled to multiple angle laser light scattering (SEC-MALLS). As previously observed^**21**,**23**^, AncCDT-5 eluted from the SEC column primarily as a monomer when injected onto the column at 8 mg/mL, while PaCDT had an elution volume consistent with its being a trimer; both of which are consistent with the crystal structures of these protein (**Figure 2**, Supplementary Figure 1, Supplementary Figure 2). The AncCDT-5 variants containing all PaCDT-interface residues (A5+CDT_interface_ and A5+CDT_interface_ΔG216) eluted with the estimated mass of trimers in solution, even at concentrations as low as 0.1 mg/mL (**Figure 2**, Supplementary Figure 1, Supplementary Figure 2). This confirms that mutations remote from the trimer interface observed in PaCDT were not required for trimerization (A5+CDT_interface_ΔG216 and PaCDT are separated by 79 substitutions remote from the trimer interface as well as the presence of the C-terminal extension in PaCDT).

The stepwise introduction of PaCDT trimer-interface residues into AncCDT-5 resulted in a switch between oligomeric states from monomer to trimer (**Figure 2, Supplementary Figure 1**). The “early” variants A5+P94L, A5+H206F, and A5+P94L+H206F (hereafter “A5.1”), and A5.1+T108R were unambiguously monomers in solution even when injected onto the column at 8 mg/mL. Variants with interfaces more similar to PaCDT, including A5.1+“YER”+T108R (i.e., 2 substitutions, Q230I+K98R, away from the trimeric A5+CDT_interface_) and A5.1+“YER”+T108R+K98R (1 substitution, Q230I, away from the trimeric A5+CDT_interface_ variant) both eluted as higher oligomeric species. These variants eluted later than the trimeric species and light-scattering was consistent with them being dimers (**Supplementary Figure 1**).

As mentioned above, some of the intermediate variants we tested contained interface residues not found in either AncCDT-5 or PaCDT; these residues (Phe101 and Val218) are found in extant proteins that, according to the original phylogenetic analysis,^21^ are descendants of AncCDT-5. A5.1+P218V remained monomeric in solution even at high concentrations (**Figure 2, Supplementary Figure 1**). A5.1+D101F also appeared to be mostly monomeric, although light-scattering did indicate a shift towards higher oligomeric forms. The combination of these two substitutions led to variant A5.1+D101F+P218V, which forms higher oligomeric states when injected onto the column at 8 mg/mL (**Figure 2, Supplementary Figure 1**). The same variant elutes as a monomer when injected onto the SEC column at 0.1 mg/mL (**Supplementary Figure 2**), indicating concentration-dependent oligomerization.

### AncCDT-5 variants can sample dimeric forms

It was notable that some of the AncCDT-5 variants, including A5.1+D101F+P218V, A5.1+“YER”+T108R and A5.1+“YER”+T108R+K98R, eluted from the column with elution volumes and MALLS scattering consistent with the presence of dimeric structures at high concentrations. A small, minor peak in the chromatogram of some monomeric variants, including AncCDT-5, is also indicative of a small proportion of dimer being present in these samples. This dimer peak is more prevalent in the alternate reconstruction of AncCDT-5, AncCDT-5(WAG),^21^ which clearly samples both monomeric and dimeric forms (**Supplementary Figure 3**).

Considering the AncCDT-5 variants only differ at positions on the PaCDT trimer interface, it is likely that any dimeric forms are asymmetric dimers that share the same interface as found between chains in the PaCDT trimer. Indeed, an AlphaFold2 structure of A5.1+D101F+P218V shows that the most likely dimeric form is an asymmetric dimer that resembles the trimeric form with one of the chains missing (**Figure 2D**). Variant A5.1+D101F+P218V (which is mostly dimer) and A5+CDT_interface_ (mostly trimer) differ at several key residues near the apex of the complex, highlighting the important role these positions have in determining the oligomeric state of the protein.

### Quaternary structure plasticity in extant homologs of PaCDT

In addition to the above variants, we also considered the oligomeric states of several extant homologs of PaCDT. In our previous work, we found that Pu1068 (a non-dehydratase homolog of PaCDT from *Candidatus* Pelagibacter ubique that is a descendant of the non-catalytic AncCDT-2) is monomeric.^21^ On the other hand, His-tagged Ws0279, a more distantly-related L-lysine binding protein (26% sequence identity with PaCDT)^21^, is a trimer in solution and crystals (**Supplementary Figure 4**). Finally, the extant homolog Ea1174 elutes as a monomer (**Supplementary Figure 5**) but has dehydratase activity comparable to PaCDT in complementation assays.^21^ Thus, in the same manner that the point mutants of AncCDT-5 easily perturb the oligomeric equilibrium, extant proteins throughout this family appear to regularly interchange between different oligomeric states throughout the phylogeny.

### Formation of the trimer corresponds with an increase in substrate affinity

The dehydratase activity of key trimer-interface variants was assessed to determine whether the formation of a higher-order oligomeric state (e.g., dimers or trimers) was associated with an increase in dehydratase activity. Similar to our previous work^23^, we noted a >50-fold increase in the catalytic efficiency between AncCDT-5 and PaCDT (**Table I, Supplementary Figure 6**). None of the variants exhibited significant changes in terms of their turnover rates (*k*_cat_), which were close to that of AncCDT-5. In contrast, we observe a significant reduction (∼7-fold) in the *K*_M_ of these variants along the trajectory, with A5+CDT_interface_ exhibiting a *K*_M_ of ∼80 µM in comparison to AncCDT-5, which has a *K*_M_ of ∼570 µM. This is particularly important in terms of the selective benefit as the physiological concentration of the substrates are likely to be in the low micromolar range (PaCDT has a *K*_M_ of ∼27 µM, for example, and the concentrations of molecules in this biosynthetic pathway are typically ∼14–18 μM in Gram-negative bacteria^24^).

**Table I.** Prephenate dehydratase activity of the variants. Values are mean ± standard error of fit (n=2-3).

| | $K_M$ ($\mu M$) | $k_{cat}$ ($s^{-1}$) | $k_{cat}/K_M$ ($M^{-1}s^{-1}$) |
| --- | --- | --- | --- |
| **A5** | 570 ± 121 | 9.11 ± 0.2 | (1.60 ± 0.38) $\times 10^4$ |
| A5.1 | 570 ± 68.6 | 10.3 ± 0.59 | (1.81 ± 0.24) $\times 10^4$ |
| A5.1+D101F | 169 ± 22.9 | 10.5 ± 0.47 | (6.22 ± 0.88) $\times 10^4$ |
| A5.1+P217V | 131 ± 37.6 | 5.99 ± 0.53 | (4.58 ± 1.38) $\times 10^4$ |
| A5.1+T108R | 178 ± 27.1 | 8.45 ± 0.43 | (4.75 ± 0.76) $\times 10^4$ |
| A5.1+D101F+P217V | 509 ± 146 | 6.90 ± 0.91 | (1.36 ± 0.43) $\times 10^4$ |
| A5.1+YER+T108R | 357 ± 83.8 | 4.88 ± 0.47 | (1.37 ± 0.35) $\times 10^4$ |
| A5.1+YER+T108R+K98R | 376 ± 24.9 | 3.93 ± 0.11 | (1.04 ± 0.07) $\times 10^4$ |
| A5.1+YER+T108R+Q229I | 203 ± 19.9 | 12.5 ± 0.43 | (6.17 ± 0.64) $\times 10^4$ |
| A5+CDT<sub>int</sub> | 81.2 ± 26.3 | 4.67 ± 0.42 | (5.75 ± 1.93) $\times 10^4$ |
| **PaCDT** | 26.7 ± 7.7 | 25.1 ± 1.60 | (94.0 ± 27.6) $\times 10^4$ |

We also tested whether trimerization was associated with an increase in the thermostability of the proteins; we reasoned that the formation of the trimer may have enhanced the thermostability, thereby providing an additional adaptive advantage. However, there was no clear correlation between the thermostability of the proteins and their oligomeric state (**Supplementary Figure 7, Supplementary Table II**). PaCDT had the lowest thermostability with a T_m_ of below 70 °C while AncCDT-5 was the most stable (T_m_ of ∼77 °C); the introduction of PaCDT trimer-interface substitutions to AncCDT-5 led to a decrease in thermostability of these variants, approximating that of PaCDT.

Together, our results suggest (i) that the oligomeric state of PaCDT is evolutionarily-metastable, with the equilibrium being easily perturbed between monomer, dimer and trimer states by mutation and (ii) that the oligomerization affects catalytic activity (*K*_M_ in particular) but not thermostability.

### The role of the C-terminal region in the activity of PaCDT

While trimerization could account for a notable decrease in *K*_M_ between AncCDT-5 and PaCDT, it could not fully account for the improved kinetic parameters of PaCDT relative to AncCDT-5. In particular, the trimeric AncCDT-5 variants, including A5+CDT_interface_, still have *k*_cat_ values that are significantly lower than that of PaCDT. Since trimerization did not fully account for the shift in kinetic properties observed between AncCDT-5 and PaCDT, we turned our attention to other structural differences between AncCDT-5 and PaCDT.

PaCDT has a nine amino acid extension at its C-terminus compared with AncCDT-5 (**Figure 3**); differences in this region have been associated with functional differences in other periplasmic binding proteins.^25^ To test the effect of this region, truncation of the C-terminal extension was performed in PaCDT, yielding a variant (PaCDTΔC) that retained a trimeric oligomeric state (**Supplementary Figure 8**). The effect on the kinetic parameters was greatest in terms of *k*_cat_, which fell to the same rate as AncCDT-5; this is notable because the trimerization of AncCDT-5 had no effect on *k*_cat_. In contrast, the *K*_M_ increased somewhat, but remained significantly lower than the *K*_M_ of AncCDT-5 (147 ± 48.8 μM vs 570 ± 9.1 μM). Thus, the C-terminal extension alone can account for most of the difference in *k*_cat_ between trimeric AncCDT-5 and trimeric PaCDT, but the trimeric truncated version retained lower *K*_M_. That is, two variants (A5+CDT_interface_ and PaCDTΔC) that are both trimeric and lack the C-terminal extension, but otherwise differ at 80 positions, both display substantially lower *K*_M_ values than monomeric AncCDT-5.

**Figure 3.**
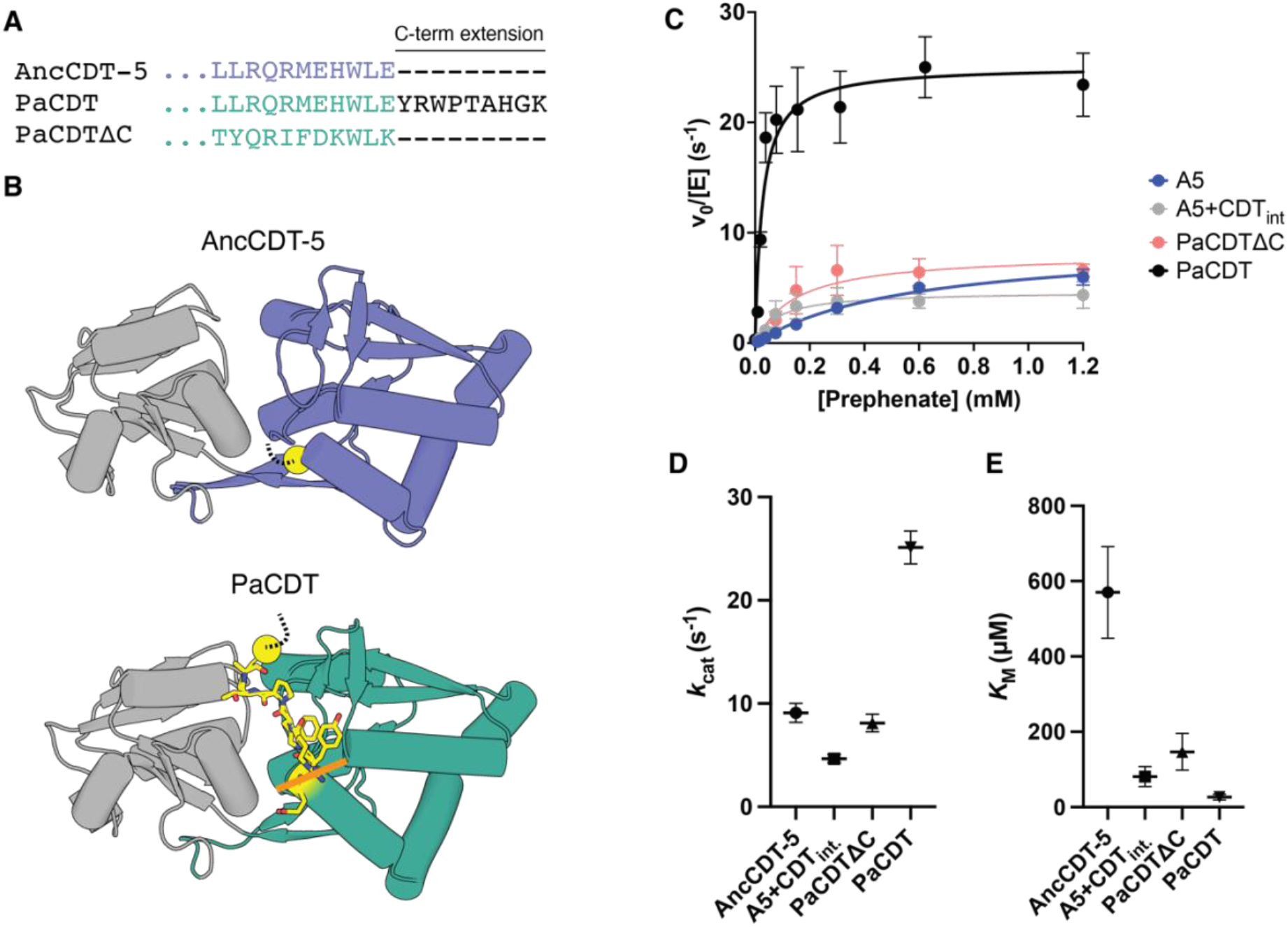
The prephenate dehydratase activity of PaCDT with the C-terminal extension truncated. **(A)** Alignment of C-terminal region of AncCDT-5, PaCDT and PaCDTΔC showing the C-terminal extension in PaCDT. **(B)** Crystal structures of AncCDT-5 (PDB 6WUP, final Glu residue is not resolved but shown as a dotted line) and PaCDT (PDB 6BQE, final two residues not resolved in crystal structures shown as dotted line), show the position of the C-terminal extension (yellow) in PaCDT and rough position of truncation (orange line). **(C)** Truncation of the C-terminal extension in PaCDT results in a protein that has prephenate dehydratase activity comparable to AncCDT-5 and A5+CDT_interface_. Data is mean ± SEM (n ≥ 2). Data for AncCDT-5, A5+CDT_interface_ and PaCDT is the same as is shown in **Table I. (D)** *k*_cat_ values as mean ± SEM (n ≥ 2). **(E)** *K*_M_ values as mean ± SEM (n ≥ 2).

### Trimerization and the C-terminal extension affect the conformational sampling of PaCDT

In our previous work, we found that PaCDT sampled the catalytically relevant closed conformation more often compared with the ancestral sequences (which predominantly sampled states in which the two subdomains were far apart).^21,23^ This provides a plausible explanation for the difference in kinetic parameters: AncCDT-5, which mostly samples a “wide-open” state that requires significant reorganization to attain a Michaelis complex has a much higher *K*_M_ and lower catalytic turnover than the extant PaCDT, which samples compact states around the Michaelis state. We speculated that trimerization or the addition of the C-terminal extension may have altered the open-closed conformational sampling in PaCDT by limiting the range of motion of the smaller domains and that this is what caused the shift in the catalytic parameters.

As such, we used molecular dynamics (MD) simulations to probe whether either trimerization or the C-terminal extension could have affected the open-closed conformational sampling of these enzymes to provide a molecular-level rationalization of the kinetic results. Our previous work on PaCDT and its ancestors demonstrated that MD simulations were consistent with experimental measurement of conformational sampling using double electron–electron resonance (DEER) spectroscopy distance measurements using unnatural amino acid:lanthanide labels.^23^ Accordingly, here we ran additional MD simulations of PaCDT both with and without the C-terminal extension and compared these to simulations of monomeric AncCDT-5. As a simple proxy for the extent of the “openness” of the protein and the pre-organization of active site residues, we monitored how the distance between the alpha carbons of the catalytic residue Glu184 on the small domain (due to different numbering conventions, this is equivalent to the catalytic residue Glu173 mentioned in our previous work^21,23^) and Tyr33 on the large domain (equivalent to Tyr22 in previous work) changed during the course of the simulations (**Figure 4A**).

Simulations initiated from structures based on the crystal structure of monomeric AncCDT-5 were consistent with our previous work^23^, showing a protein that regularly samples wide-open states (in which the small domain completely twists away from the large domain, resulting in ∼3 nm between the two key residues) (**Figure 4B**, **Supplementary Figure 9**). When these wide-open states of AncCDT-5 are overlaid on the trimeric structure of PaCDT, it is clear that in a trimeric arrangement it would not be possible for more than one of the three chains to simultaneously sample this conformation since the small domains of each chain would clash (**Figure 4C**). Consistent with this, simulations initiated from PaCDT in which the C-terminal extension had been removed, which experimentally retained a trimeric structure and displayed similar *k*_cat_ but significantly lower *K*_M_ values than AncCDT-5, showed reduced sampling of the wide-open, twisted state, with more regular sampling of semi-open (∼1.5 nm) and compact conformations (**Figure 4D**). Importantly, the arrangement of chains in the trimer prevented the chains opening to the same extent as observed in the AncCDT-5 simulations, and also prevented all three chains from opening simultaneously (i.e., at least one chain was always in a more closed, catalytically relevant conformation; <1.5 nm; **Figure 4E**, Supplementary Figure 9). Finally, simulations of the full length PaCDT trimer (i.e. with the C-terminal extension) showed that it sampled a relatively narrow distribution of closed states for the majority of the simulations, with limited sampling of a semi-open state (∼1.5 nm) that would allow for substrate-binding and release and no sampling of the wide-open state (i.e. Glu184–Tyr33 distances remained below 2 nm; **Figure 4F,G**), consistent with previous work.^23^ Visual inspection of the trajectories revealed frequent bridging interactions between the small and large domains that were mediated by the C-terminal region. Thus, the formation of a trimer and the subsequent addition of a C-terminal extension progressively reshapes the conformational landscape of CDT to freeze out non-catalytic conformations.

**Figure 4.**
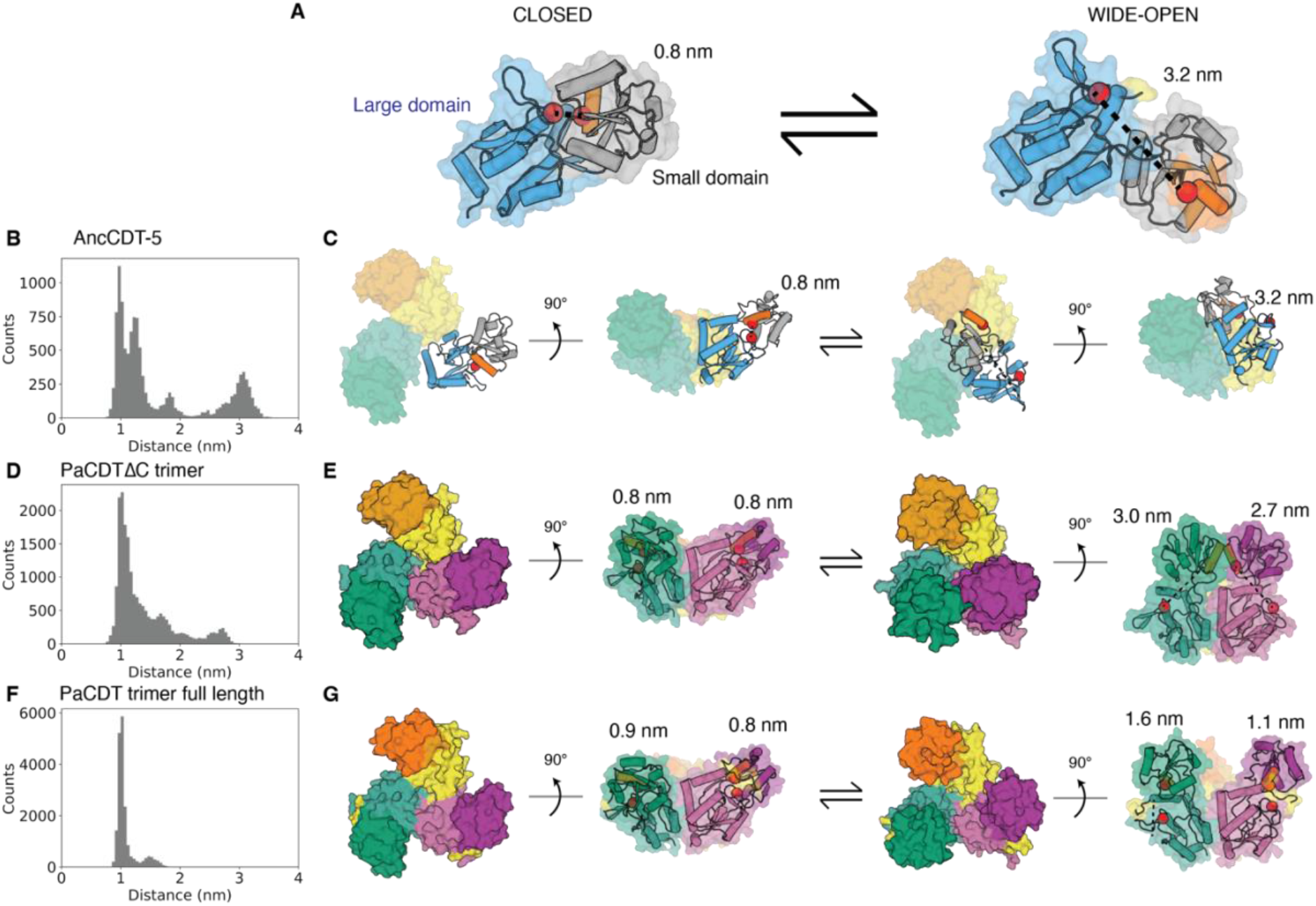
Molecular dynamics simulations of AncCDT-5, PaCDTΔC and PaCDT. **(A)** Monomeric AncCDT-5 sampled a range of conformations ranging from a more closed state (left) in which the catalytic residues are close together to a distorted “wide-open” state (right) in which the small domain is twisted relative to the large domain catalytic residues are far apart (> 2.5 nm). During simulations we monitored the distance between alpha-carbons of Glu184 and Tyr33 (red spheres). The position of the C-terminus is highlighted in yellow. The orientation of the helix on which Glu184 is found is highlighted in orange, and the small and large subdomains are shaded grey and blue, respectively. **(B)** Histogram showing the distribution of Glu184–Tyr33 distances during replicate simulations of AncCDT-5 (the first 50 ns of each simulation was omitted from histogram data). **(C)** Snapshots corresponding to closed (left) and wide-open (right) states of AncCDT-5 (cartoon representation) are mapped onto the trimeric structure of PaCDT, showing that sampling of the wide-open state would be restricted in a trimeric arrangement even when the other two chains (yellow and green surfaces) are in a closed conformation. **(D)** Histogram showing the distribution of Glu184–Tyr33 distances during replicate simulations of PaCDTΔC. **(E)** In simulations of PaCDTΔC, the small domains did not interact when they were all in a more closed state (left), but steric clashes between the small subunits of neighboring open chains corresponded to reduced sampling of the wide-open states (right). **(F)** Histogram showing the distribution of Glu184–Tyr33 distances during replicate simulations of full-length PaCDT. **(G)** Full-length PaCDT sampled a much more compact state (left), with the additional C-terminal residues keeping catalytic residues near even in the most open of the MD snapshots (right).

## Discussion

While the majority of experimentally characterized proteins with the periplasmic binding protein-like II fold are monomeric^26^ (with a few notable examples of dimeric assemblies^11,27–29^), in this work we show that PaCDT homologs and ancestral states interconvert easily between a range of oligomeric structures including monomers, dimers and trimers. Indeed, the oligomeric state of the AncCDT-5 variants can be readily shifted with as few as one or two substitutions at the trimer-forming interfaces, suggesting that different oligomeric states could be accessed easily via neutral genetic drift. There also appears to be significant quaternary structure plasticity amongst extant PaCDT homologs including Ws0279, Pu1068 and Ea1174. As such, the oligomeric states that PaCDT variants can sample appear to be evolutionarily metastable in the sense that the oligomeric equilibrium can be readily switched via a small number of mutations. This is consistent with observations from other studies that have traced the evolution of oligomerization. For example, as few as two mutations were required to switch between dimeric into a tetrameric states during the evolution of vertebrate haemoglobin^8^. Additionally, quaternary structure plasticity has been attributed to functional diversification in other protein families, including RuBisCO ^6^ and methionine S-adenosyltransferase^20^.

While other studies have retraced historical dimerization events (which typically involve the evolution of a single surface), the work presented here shows that cyclic oligomers that involve heterologous interfaces can also be readily accessible via a small number of mutations when these are introduced into an appropriate genetic background. This is notable considering that for cyclic complexes to emerge it requires the evolution and optimization of two distinct complementary interfaces. Not only this, but the resulting geometry must also be such that it can accommodate packing of multiple chains around the point of symmetry. Indeed, these factors contribute to the rarity of trimers amongst natural proteins.^30^ Here, we have provided structural rationalization for the emergence of the PaCDT and highlighted the importance of residues near the apex of the trimer in determining the oligomeric state of the protein. As such, this work furthers our understanding of the emergence of cyclic protein complexes which are important targets for both drug design^31,32^ and protein engineering.^15,19^

The effect of oligomerization on enzyme activity is often somewhat cryptic as it is often unclear how an interface remote from an active site can affect activity. Moreover, it is difficult to test with extant proteins because of epistasis and the ratchet-like nature of evolution^12,33,34^; if an interface is disrupted, how can we be sure the effects are from the loss of the oligomeric structure and not other structural changes? Constructive biochemistry approaches such as ancestral sequence reconstruction are useful in this context, as they allow us to see the benefits of oligomerization in the “forward-direction”. In this case, we can see that formation of a trimer generates a more enzyme-like conformational landscape that favors the formation of the enzyme Michaelis complex and decreases *K*_M_ to physiologically relevant concentrations.

It is well established that historical events (including oligomerization) shape future evolutionary trajectories, even if the initial event is neutral or near-neutral in terms of providing a selective advantage.^35–38^ Genetic variation within protein families can lead to (often cryptic) variations in protein properties that influence the evolvability of a protein and subsequent evolutionary trajectories. Such substitutions and properties can become entrenched by subsequent mutations that prevent reversion to the ancestral state.^14^ While the formation of the trimer provided an immediate enhancement in *K*_M_ during the evolution of PaCDT, trimerization would have also determined which future evolutionary trajectories were accessible and thus shaped any subsequent optimization of the enzyme; this may have included modifications in the C-terminal region that we showed contributed to an increase in the catalytic turn-over of the enzyme. At the same time, it is conceivable that efficient dehydratase activity may have been achievable without trimerization via other mutational routes; indeed, the modern enzyme Ea1174 seems to be an efficient dehydratase despite being a monomer. Further, significant variation in the C-terminal regions of CDT homologs suggest that this region may be important for additional fine-tuning of activity amongst different species.

## Conclusion

Considering the prevalence and biological importance of homo-oligomers, understanding the drivers and consequences of oligomerization is important to understand protein biology and enable better protein design. Here, we show that the oligomeric state of PaCDT ancestors and homologs can be ready perturbed through a minimal number of substitutions at the interface, and that this can have significant effects on activity by altering the conformational sampling of the proteins. This suggests that new oligomeric states (such as the trimeric form seen in PaCDT and Ws0279) may have initially emerged via the accumulation of relatively neutral mutations and subsequently entrenched the higher order oligomeric state.

## Materials and Methods

### Design of interface mutants

The sequences of trimer-interface variants were based on the sequence of AncCDT-5, an inferred ancestral sequence from our previous work.^21,23^ Unless otherwise stated, residue numbering used in this study is based on the numbering used in the crystal structure of AncCDT-5 (PDB 6WUP); as such, the numbering convention used here differs from the residue numbering convention used in our previous work.^21,23^

The trimer-interface-forming residues were identified by analysing crystal structures of the trimeric PaCDT (PDB IDs 5HPQ, 6BQE) and using the InterfaceResidues script in PyMOL (https://pymolwiki.org/index.php/InterfaceResidues). Sequence and structural alignments between AncCDT-5 and PaCDT were then used to identify positions that differed between AncCDT-5 and PaCDT at these trimer-forming interface positions; this guided the design of AncCDT-5 variants that differed only at these trimer-interface positions. In order to roughly reflect the historical order in which trimer-forming residues were introduced, we also considered the sequences of PaCDT homologs that were identified to be decedents of AncCDT-5 in our previous work.^21^ In addition, PAML^39^ was used to infer the maximum-likelihood amino acid sequences at internal nodes between AncCDT-5 and PaCDT in the LG-inferred phylogenetic tree described in our previous work.^21^ The presence or absence of gaps in the inferred ancestral proteins was decided based on parsimony and manual inspection of the multiple-sequence alignment. In addition to a variant of AncCDT-5 that had all the interface positions mutated to their corresponding PaCDT residues (“A5+CDT_interface_”), we also generated a similar protein with Gly216 removed (“A5+CDT_interface_ΔG216”). We also constructed AncCDT-5 variants that introduced other residues found in other PaCDT homologs at the positions identified at the apex of the PaCDT trimer, including Val218 and Phe101. Sequences were back-translated, and the resulting DNA fragments encoding these variants were codon-optimized for expression in *E. coli*, synthesized and cloned into the pET28a vector (between BamHI and XhoI restriction sites) by Twist Bioscience. Variants A5.1+D101F, A5.1+P217V, A5.1+T108R and A5.1+YER+T108R+Q229I were generated using mutagenesis primers (**Supplementary Table IV**) using a standard Gibson Assembly protocol.^40^ pET28a vectors containing the genes encoding AncCDT-5^21^, PaCDT (UniProt: Q01269; residues 26–268) and the C-terminal-truncated PaCDT (UniProt: Q01269; residues 26 to 259) were also obtained. The pET28a vector adds a pET28a-encoded N-terminal hexahistadine tag, thrombin protease cleavage sequence and T7 tag to the variant sequences. While the resulting protein sequences used in this study are similar to those encoded in pDOTS7 vectors used in our previous work^21,23^, they do have slightly different N-terminal tags (e.g. additional T7 tag) and the pET28a-encoded genes also omit the additional C-terminal leucine residue found in the pDOTS7-based constructs as a consequence of the Golden-Gate assembly method that was used. For oligomeric characterization of Ws0279 (UniProt: Q7MAG0; residues 24–258) and Ea1174 (UniProt: K0ABP5; residues 31–268), pDOTS7 vectors encoding the proteins were obtained from our previous work.^21^ Similarly, AncCDT-5(WAG) was characterized by expressing it from a pDOTS7 vectors obtained from our previous work.

### Expression and purification of protein variants

#### Protein Expression

In most cases, variants were expressed in BL21(DE3) cells (NEB): plasmids were incorporated via electroporation, followed by recovery in LB media for 1 h and incubating at 37 °C on LB agar plates supplemented with 50 µg/mL kanamycin. The following day, single colonies were used to inoculate 5 mL starter cultures (LB media + 50 µg/mL kanamycin), which were incubated for 5-6 h at 37 °C with shaking (∼180 rpm). These were then used to inoculate 1 L LB + 50 µg/mL kanamycin cultures in Thomson Ultra Yield Flasks. Cultures were grown at 37 °C with shaking (180 rpm) until OD_600_ reached approximately 0.6 before being induced with 0.5 mM β-d-1-isopropylthiogalactopyranoside (IPTG). Following induction, cultures were grown overnight at 30 °C with shaking at 180 rpm. Cell pellets were collected by centrifugation and stored at −20 °C. His-tagged Ws0279, Ea1174 and AncCDT-5(WAG) were expressed in BL21(DE3) cells (NEB): transformed cells were grown in LB media supplemented with 100 mg/L ampicillin to OD_600_ ∼0.7 at 37°C, induced with 1 mM IPTG, and incubated for a further 20 h at 37°C.

#### Purification by Immobilized Metal Affinity Chromatography (IMAC)

All variants were initially purified from cell lysate using a standard immobilized-metal affinity chromatography (IMAC) approach. Briefly, cell pellets were thawed and resuspended in binding buffer (50 mM NaH_2_PO_4_, 500 mM NaCl, 20 mM imidazole, pH 7.4) and lyzed by sonication on ice (50% power, two rounds of 6 minutes on, 6 minutes off). Lyzed cells were fractionated by ultracentrifugation (9000 × g for 40 minutes at 4 °C). The supernatant was filtered through a 0.45 μm syringe filter and loaded onto a 5 mL HisTrap HP column (GE Healthcare) equilibrated in binding buffer. The column was washed with 30 mL of binding buffer followed by 25 mL of wash buffer (50 mM NaH_2_PO_4_, 500 mM NaCl, 40 mM imidazole, pH 7.4). The His-tagged protein was eluted in elution buffer (50 mM Na_2_HPO_4_, 500 mM NaCl, 500 mM imidazole, pH 7.4). Fractions containing protein were pooled. Samples collected from IMAC were exchanged into size exclusion chromatography (SEC) buffer (20 mM Na_2_HPO_4_, 150 mM NaCl, pH 7.4) using either a HiPrep 26/10 desalting column (GE Healthcare) or through multiple rounds of concentration and dilution using an Amicon Ultra-15 filter unit with a 10 kDa molecular weight cut-off. Protein purity was determined through sodium dodecyl sulphate–polyacrylamide gel electrophoresis (SDS-PAGE). The concentrations of purified protein samples were determined by measuring A_280_ using a NanoDrop One (Thermo Scientific) and molar extinction coefficients calculated from ProtParam (http://expasy.org/tools/protparam.html).

### Protein characterization

#### Size exclusion chromatography (SEC) and comparison with protein standards

SEC was performed at 4 °C on an ÄKTA Pure. Samples were injected onto either a HiLoad 16/600 Superdex 200 column (GE Healthcare) or a Superdex 200 10/300 GL size-exclusion column (GE Healthcare) and purified in SEC buffer. Protein standards (GE Healthcare Gel Filtration Calibration Kits) were used for comparison of elution volumes and estimation of theoretical molecular masses (**Supplementary Figure 10, Supplementary Figure 11**).

#### SEC coupled with multiple angle light scattering (SEC-MALLS)

Samples (100 μL) of each protein were loaded at 5-8 mg mL^−1^ onto a pre-equilibrated Superdex 200 10/300 GL size-exclusion column (GE Healthcare) attached to multi-angle light scattering (DAWN HELEOS 8; Wyatt Technologies) and refractive index detection (Optilab rEX; Wyatt Technologies) units. A flow rate of 0.5 mL min^−1^ was used. The multi-angle detectors were normalized using monomeric bovine serum albumin (Sigma, A1900). A dn/dc value of 0.186 g^−1^ was used for each sample. The data were processed using ASTRA 5.3.4 (Wyatt Technologies). Unless otherwise stated, data were collected from a single experiment for each variant (n = 1).

#### Differential Scanning Fluorimetry (DSF) - Thermostability measurements

Differential scanning fluorimetry (DSF) was performed on a QuantStudio 3 Real-Time-PCR-System (Thermo Fisher). In a 96 well PCR plate, reaction mixtures containing approximately 5 μM protein in DSF buffer (50 mM Na_2_HPO_4_, 150 mM NaCl, pH 7.4) and 1x Protein Thermal Shift Dye (Thermo Fisher) were heated from 20 °C to 99 °C, while monitoring emission at 623 nm with excitation at 580 nm. Measurements were completed as technical triplicates, with melting temperatures (T_m_s) calculated determined from the steepest part of the curve (i.e., peaks in the derivative plot) in the Protein Thermal Shift Software v1.4 (Thermo Fisher).

#### Prephenate dehydratase activity assays

Sodium prephenate was prepared by mixing 40 mM barium chorismite (Sigma) in water, followed by addition of an equimolar amount of Na_2_SO_4_. To this an equal volume of 100 mM Na_2_HPO_4_ was added causing precipitation of barium sulfate which was subsequently removed through centrifugation. The resulting sodium chorismite solution was heated at 70 °C for 1 h to generate sodium prephenate. The concentration of prephenate was determined through acid conversion on prephenate to phenylpyruvate using 0.5 M HCl over a 15-minute time-course, followed by the addition of NaOH and measurement of absorbance at 320 nm using the extinction coefficient for phenylpyruvate (17,500 M^−1^cm^−1^).

PDT activity was determined by using a discontinuous colorimetric assay to monitor the production of phenylpyruvate. Enzyme solutions were prepared in SEC buffer to between 100-700 nM. Sodium prephenate was diluted to 1.4 mM in 50 mM Na_2_HPO_4_ and a series of seven 1-in-2 dilutions was created. To initiate the assay, 100 μL of the enzyme solution (final concentration of 100 nM) was mixed with 600 μL of substrate at 25 °C. Every ∼30 seconds, the absorbance of a 100 μL aliquot of enzyme-substrate solution was measured at 320 nm following the addition of 100 μL of 2 M NaOH. No-enzyme controls were performed; there was negligible product formation during the time-course of the assays (∼approximately 3 mins). The concentration of phenylpyruvate was calculated assuming a pathlength of 0.5 cm, using the previously reported extinction coefficient (17,500 M^−1^cm^−1^), and taking into account dilution factors. Data was plotted and analyzed using GraphPad Prism (version 9) using the Michaelis-Menten model.

### Modelling using ColabFold

The ColabFold: AlphaFold2 using MMseqs Google Colab notebook^41,42^ was used to generate the model of dimeric A5.1+D101F+P217V: default options were used (with the addition of an amber minimization step).

### Molecular dynamics simulations

Molecular dynamics (MD) simulations were performed using Gromacs 2021.2^43^ and the Charmm36m force field (charmm36-feb2021.ff)^44^. Several systems were prepared (see **Supplementary Table III** for details). In each case, protein structures were obtained from the PDB and then submitted through the PDB REDO server.^45^ Small molecules (including waters and acetate/HEPES) were removed. Protein structures were further prepared using the Schrodinger Protein Preparation Tool (Schrodinger 2020-4), including removal of unwanted chains, ionization of residues (pH 7.4), optimization of H-bonds and energy minimization. In cases where residues not observed in the crystal structure were modelled, the Schrödinger modelling tools were used to model the C-terminal extensions followed by local minimization in Maestro. Protein chain termini were capped in Gromacs. Each protein was solvated in a rhombic dodecahedron with SPC water molecules, such that the minimal distance of the protein to the periodic boundary was 14 Å, and an appropriate number of ions were added to neutralize the system (see **Supplementary Table III**). Energy minimization was performed on each system using the steepest descent algorithm, followed by a 100 ps isothermal-isochoric ensemble (NVT; 300 K) simulation with harmonic position restrains on the protein’s heavy atoms. This was followed by a 100 ps NPT simulation with harmonic position restrains on the protein’s heavy atoms. Production MD simulation runs were maintained at 300 K using a V-rescale thermostat (τT = 1 ps) and 1 bar using the C-rescale barostat (τp = 2.0 ps, compressibility = 4.5 × 10^−5^ bar^−1^). Hydrogen bonds were constrained using the LINCS algorithm. The cut-off for short-range electrostatics was 1.2 Å. For each system, at least four random-seed replicates were performed. All systems were run for 500 ns per replicate. For analysis, frames corresponding to every 200 ps (i.e., 0.2 ns) were extracted (using gmx trjconv). RMSD and distance measurements were calculated using gromacs tools (gmx rms and gmx distance). Data was plotted using matplotlib.

## Supplementary Material

Supplementary Information File – contains supplementary figures and tables.

## Data availability

All relevant data and scripts have been deposited at 10.5281/zenodo.6894455. All other data are available from authors upon request.

## Acknowledgements

This research was conducted by the Australian Research Council Centre of Excellence in Synthetic Biology (project number CE200100029) and funded by the Australian Government. CJJ also acknowledges support from the ARC Centre of Excellence for Innovations in Peptide and Protein Science (CIPPS). BEC was supported by a JSPS Postdoctoral Fellowship for Overseas Researchers and KAKENHI Grant-in-Aid for Scientific Research (20F20705). This research/project was undertaken with the assistance of resources and services from the National Computational Infrastructure (NCI), which is supported by the Australian Government.

## References

1. Levy ED, Teichmann SA. Structural, evolutionary, and assembly principles of protein oligomerization. Progress in Molecular Biology and Translational Science. 2013;117:25–51. doi:10.1016/b978-0-12-386931-9.00002-7

2. Marsh JA, Rees HA, Ahnert SE, Teichmann SA. Structural and evolutionary versatility in protein complexes with uneven stoichiometry. Nature Communications. 2015;6(1):6394–6394. doi:10.1038/ncomms7394

3. Levy ED, Pereira-Leal JB, José B. Pereira-Leal, Pereira-Leal JB, Chothia C, Teichmann SA. 3D complex: a structural classification of protein complexes. PLOS Computational Biology. 2005;2(11). doi:10.1371/journal.pcbi.0020155

4. Kühner S, van Noort V, Betts MJ, Leo-Macias A, Batisse C, Rode M, Yamada T, Maier T, Bader SJ, Beltran-Alvarez P, et al. Proteome Organization in a Genome-Reduced Bacterium. Science. 2009;326(5957):1235–1240. doi:10.1126/science.1176343

5. H. K. Schachman, Schachman HK. From allostery to mutagenesis: 20 years with aspartate transcarbamoylase. Biochemical Society Transactions. 1987;15(4):772– 775. doi:10.1042/bst0150772

6. AK Liu, JH Pereira, AJ Kehl, DJ Rosenberg, DJ Orr, SKS Chu, DM Banda, M Hammel, PD Adams, JB Siegel, et al. Structural plasticity enables evolution and innovation of RuBisCO assemblies. Science advances. 2022. doi:10.1126/sciadv.adc9440

7. Perica T, Marsh JA, Sousa FL, Natan E, Colwell LJ, Ahnert SE, Teichmann SA. The emergence of protein complexes: quaternary structure, dynamics and allostery. Biochemical Society Transactions. 2012;40(3):475–491. doi:10.1042/bst20120056

8. Pillai AS, Chandler SA, Liu Y, Signore AV, Cortez-Romero CR, Benesch JLP, Laganowsky A, Storz JF, Hochberg GKA, Thornton JW. Origin of complexity in haemoglobin evolution. Nature. 2020;581(7809):480–485. doi:10.1038/s41586-020-2292-y

9. Bergendahl LT, L. Therese Bergendahl, Marsh JA. Functional determinants of protein assembly into homomeric complexes. Scientific Reports. 2017;7(1):4932– 4932. doi:10.1038/s41598-017-05084-8

10. Fraser NJ, Liu J-WW, Mabbitt PD, Correy GJ, Coppin CW, Lethier M, Perugini MA, Murphy JM, Oakeshott JG, Weik M, et al. Evolution of Protein Quaternary Structure in Response to Selective Pressure for Increased Thermostability. Journal of Molecular Biology. 2016;428(11):2359–2371. doi:10.1016/j.jmb.2016.03.014

11. Ruggiero A, Dattelbaum JD, Staiano M, Berisio R, D’Auria S, Vitagliano L. A loose domain swapping organization confers a remarkable stability to the dimeric structure of the arginine binding protein from Thermotoga maritima. PLOS ONE. 2014;9(5):e96560–e96560. doi:10.1371/JOURNAL.PONE.0096560

12. Luca Schulz, Franziska L. Sendker, Hochberg GKA. Non-adaptive complexity and biochemical function. Current Opinion in Structural Biology. 2022;73:102339–102339. doi:10.1016/j.sbi.2022.102339

13. Muñoz-Gómez SA, Bilolikar G, Wideman JG, Geiler-Samerotte K. Constructive Neutral Evolution 20 Years Later. Journal of Molecular Evolution. 2021;89(3):172– 182. doi:10.1007/s00239-021-09996-y

14. Hochberg GKA, Liu Y, Marklund EG, Metzger BPH, Laganowsky A, Thornton JW. A hydrophobic ratchet entrenches molecular complexes. Nature. 2020;588(7838):503–508. doi:10.1038/s41586-020-3021-2

15. Zhu J, Avakyan N, Kakkis A, Hoffnagle AM, Han K, Li Y, Zhang Z, Tae Su Choi, Choi TS, Na Y, et al. Protein Assembly by Design. Chemical Reviews. 2021 Aug 18. doi:10.1021/acs.chemrev.1c00308

16. Ueda G, Aleksandar Antanasijevic, Aleksandar Antanasijevic, Antanasijevic A, Fallas JA, Sheffler W, Copps J, Ellis D, Geoffrey B. Hutchinson, Hutchinson GB, et al. Tailored design of protein nanoparticle scaffolds for multivalent presentation of viral glycoprotein antigens. eLife. 2020;9. doi:10.7554/elife.57659

17. Park J, Selvaraj B, McShan AC, Boyken SE, Wei KY, Oberdorfer G, DeGrado WF, Sgourakis NG, Cuneo MJ, Matthew J. Cuneo, et al. De novo design of a homo-trimeric amantadine-binding protein. eLife. 2019;8. doi:10.2210/pdb6naf/pdb

18. Boyken SE, Chen Z, Groves B, Langan RA, Oberdorfer G, Ford A, Gilmore JM, Xu C, DiMaio F, Pereira JH, et al. De novo design of protein homo-oligomers with modular hydrogen-bond network–mediated specificity. Science. 2016;352(6286):680– 687. doi:10.1126/science.aad8865

19. Fallas JA, Ueda G, Sheffler W, Vanessa Nguyen, McNamara DE, Sankaran B, Pereira JH, Fabio Parmeggiani, Parmeggiani F, Brunette TJ, et al. Computational design of self-assembling cyclic protein homo-oligomers. Nature Chemistry. 2017;9(4):353–360. doi:10.1038/nchem.2673

20. Kleiner D, Shapiro Tuchman Z, Shmulevich F, Shahar A, Zarivach R, Kosloff M, Bershtein S. Evolution of homo-oligomerization of methionine S-adenosyltransferases is replete with structure–function constrains. Protein Science. 2022 [accessed 2022 Aug 15];31(7). https://onlinelibrary.wiley.com/doi/10.1002/pro.4352. doi:10.1002/pro.4352

21. Clifton BE, Kaczmarski JA, Carr PD, Gerth ML, Tokuriki N, Jackson CJ. Evolution of cyclohexadienyl dehydratase from an ancestral solute-binding protein. Nature Chemical Biology. 2018;14(6):542–547. doi:10.1038/s41589-018-0043-2

22. Zhao GS, Xia TH, Fischer RS, Jensen RA. Cyclohexadienyl dehydratase from Pseudomonas aeruginosa. Molecular cloning of the gene and characterization of the gene product. J. Biol. Chem. 1992;267.

23. Kaczmarski JA, Mahawaththa MC, Feintuch A, Clifton BE, Adams LA, Goldfarb D, Otting G, Jackson CJ. Altered conformational sampling along an evolutionary trajectory changes the catalytic activity of an enzyme. Nature Communications. 2020;11(1):5945. doi:10.1038/s41467-020-19695-9

24. Bennett BD, Kimball EH, Gao M, Osterhout R, Van Dien SJ, Rabinowitz JD. Absolute metabolite concentrations and implied enzyme active site occupancy in Escherichia coli. Nature Chemical Biology. 2009;5(8):593–599. doi:10.1038/nchembio.186

25. Gouridis G, Muthahari YA, de Boer M, Griffith DA, Tsirigotaki A, Tassis K, Zijlstra N, Xu R, Eleftheriadis N, Sugijo Y, et al. Structural dynamics in the evolution of a bilobed protein scaffold. Proceedings of the National Academy of Sciences of the United States of America. 2021;118(49). doi:10.1073/pnas.2026165118

26. Scheepers GH, Lycklama a Nijeholt JA, Poolman B. An updated structural classification of substrate-binding proteins. FEBS Letters. 2016;590(23):4393–4401. doi:10.1002/1873-3468.12445

27. Shi R, Proteau A, Wagner J, Cui Q, Purisima EO, Matte A, Cygler M. Trapping open and closed forms of FitE: a group III periplasmic binding protein. Proteins. 2009;75(3):598–609. doi:10.1002/prot.22272

28. Li L, Ghimire-Rijal S, Lucas SL, Stanley CB, Wright E, Agarwal PK, Myles DAA, Cuneo MJ. Periplasmic Binding Protein Dimer Has a Second Allosteric Event Tied to Ligand Binding. Biochemistry. 2017;56(40):5328–5337. doi:10.1021/acs.biochem.7b00657

29. Gonin S, Arnoux P, Pierru B, Bénédicte Pierru, Pierru B, Lavergne J, Alonso B, Sabaty M, Pignol D. Crystal structures of an Extracytoplasmic Solute Receptor from a TRAP transporter in its open and closed forms reveal a helix-swapped dimer requiring a cation for α-keto acid binding. BMC Structural Biology. 2007;7(1):11–11. doi:10.1186/1472-6807-7-11

30. Lynch M. Evolutionary diversification of the multimeric states of proteins. Proceedings of the National Academy of Sciences. 2013 [accessed 2022 Aug 15];110(30). https://pnas.org/doi/full/10.1073/pnas.1310980110. doi:10.1073/pnas.1310980110

31. Bai Y, Yu Z-H, Liu S, Liu S, Zhang L, Zhang R-Y, Zeng L-F, Zhang S, Zhang ZY. Novel Anticancer Agents Based on Targeting the Trimer Interface of the PRL Phosphatase. Cancer Research. 2016;76(16):4805–4815. doi:10.1158/0008-5472.can-15-2323

32. Wrobel AG, Benton DJ, Hussain S, Harvey R, Martin SR, Martin SR, Roustan C, Rosenthal PB, Skehel JJ, Gamblin SJ. Antibody-mediated disruption of the SARS-CoV-2 spike glycoprotein. Nature Communications. 2020;11(1):5337–5337. doi:10.1038/s41467-020-19146-5

33. Ben-David M, Soskine M, Dubovetskyi A, Cherukuri K-P, Dym O, Sussman JL, Liao Q, Szeler K, Klaudia Szeler, Kamerlin SCL, et al. Enzyme Evolution: An Epistatic Ratchet versus a Smooth Reversible Transition. Molecular Biology and Evolution. 2020;37(4):1133–1147. doi:10.1093/molbev/msz298

34. Bridgham JT, Ortlund EA, Thornton JW. An epistatic ratchet constrains the direction of glucocorticoid receptor evolution. Nature. 2009;461(7263):515–519. doi:10.1038/nature08249

35. Harms MJ, Thornton JW. Historical contingency and its biophysical basis in glucocorticoid receptor evolution. Nature. 2014;512(7513):203–207. doi:10.1038/nature13410

36. Shah P, McCandlish DM, Plotkin JB. Contingency and entrenchment in protein evolution under purifying selection. Proceedings of the National Academy of Sciences. 2015;112(25):E3226–E3235. doi:10.1073/pnas.1412933112

37. Xie VC, Pu J, Metzger BP, Thornton JW, Dickinson BC. Contingency and chance erase necessity in the experimental evolution of ancestral proteins Courtier-Orgogozo V, editor. eLife. 2021;10:e67336. doi:10.7554/eLife.67336

38. Baier F, Hong N, Yang G, Pabis A, Miton CM, Barrozo A, Carr PD, Kamerlin SC, Jackson CJ, Tokuriki N. Cryptic genetic variation shapes the adaptive evolutionary potential of enzymes. eLife. 2019;8:e40789. doi:10.7554/eLife.40789

39. Yang Z. PAML 4: phylogenetic analysis by maximum likelihood. Mol. Biol. Evol. 2007;24. https://doi.org/10.1093/molbev/msm088. doi:10.1093/molbev/msm088

40. Gibson DG. Enzymatic assembly of DNA molecules up to several hundred kilobases. Nat. Methods. 2009;6. https://doi.org/10.1038/nmeth.1318. doi:10.1038/nmeth.1318

41. Mirdita M, Schütze K, Moriwaki Y, Heo L, Ovchinnikov S, Steinegger M. ColabFold: making protein folding accessible to all. Nature Methods. 2022;19(6):679–682. doi:10.1038/s41592-022-01488-1

42. Jumper J, Evans R, Pritzel A, Green T, Figurnov M, Ronneberger O, Tunyasuvunakool K, Bates R, Žídek A, Potapenko A, et al. Highly accurate protein structure prediction with AlphaFold. Nature. 2021;596(7873):583–589. doi:10.1038/s41586-021-03819-2

43. Abraham MJ, Murtola T, Schulz R, Páll S, Smith JC, Hess B, Lindahl E. GROMACS: High performance molecular simulations through multi-level parallelism from laptops to supercomputers. SoftwareX. 2015;1:19–25. doi:10.1016/j.softx.2015.06.001

44. Huang J, Rauscher S, Nawrocki G, Ran T, Feig M, de Groot BL, Grubmüller H, MacKerell AD. CHARMM36m: An improved force field for folded and intrinsically disordered proteins. Nature Methods. 2017;14(1):71–73. doi:10.1038/nmeth.4067

45. Joosten RP, Long F, Murshudov GN, Perrakis A. The PDB_REDO server for macromolecular structure model optimization. IUCrJ. 2014;1(4):213–220. doi:10.1107/S2052252514009324

