## Supplementary Information File for "The role of evolutionarily metastable oligomeric states in the optimization of catalytic activity"

Nicholas J. East - 0000-0002-4471-449X

Ben E. Clifton - 0000-0002-7469-7424

Colin J. Jackson - 0000-0001-6150-3822

Joe A. Kaczmarski - 0000-0001-8549-0122

**Affiliations**

1. ARC Centre of Excellence in Synthetic Biology, Australian National University, Canberra, Australia
2. Research School of Biology, Australian National University. R.N Robertson Building, Building 46, Biology Place, Acton, Australian Capital Territory, Australia 2601
3. ARC Centre of Excellence for Innovations in Peptide and Protein Science, Research School of Chemistry, Australian National University. Building 137, Sullivans Creek Rd, Acton, Australian Capital Territory, Australia 2601
4. Protein Engineering and Evolution Unit, Okinawa Institute of Science and Technology, 1919-1 Tancha, Onna, Okinawa, Japan 904-0412

*** Corresponding author:**

- Joe A Kaczmarski, Research School of Biology, Australian National University. R.N Robertson Building, Building 46, Biology Place, Acton, Australian Capital Territory, Australia 2601.

**Running title (50 characters or less):** Oligomerization during the evolution of PaCDT

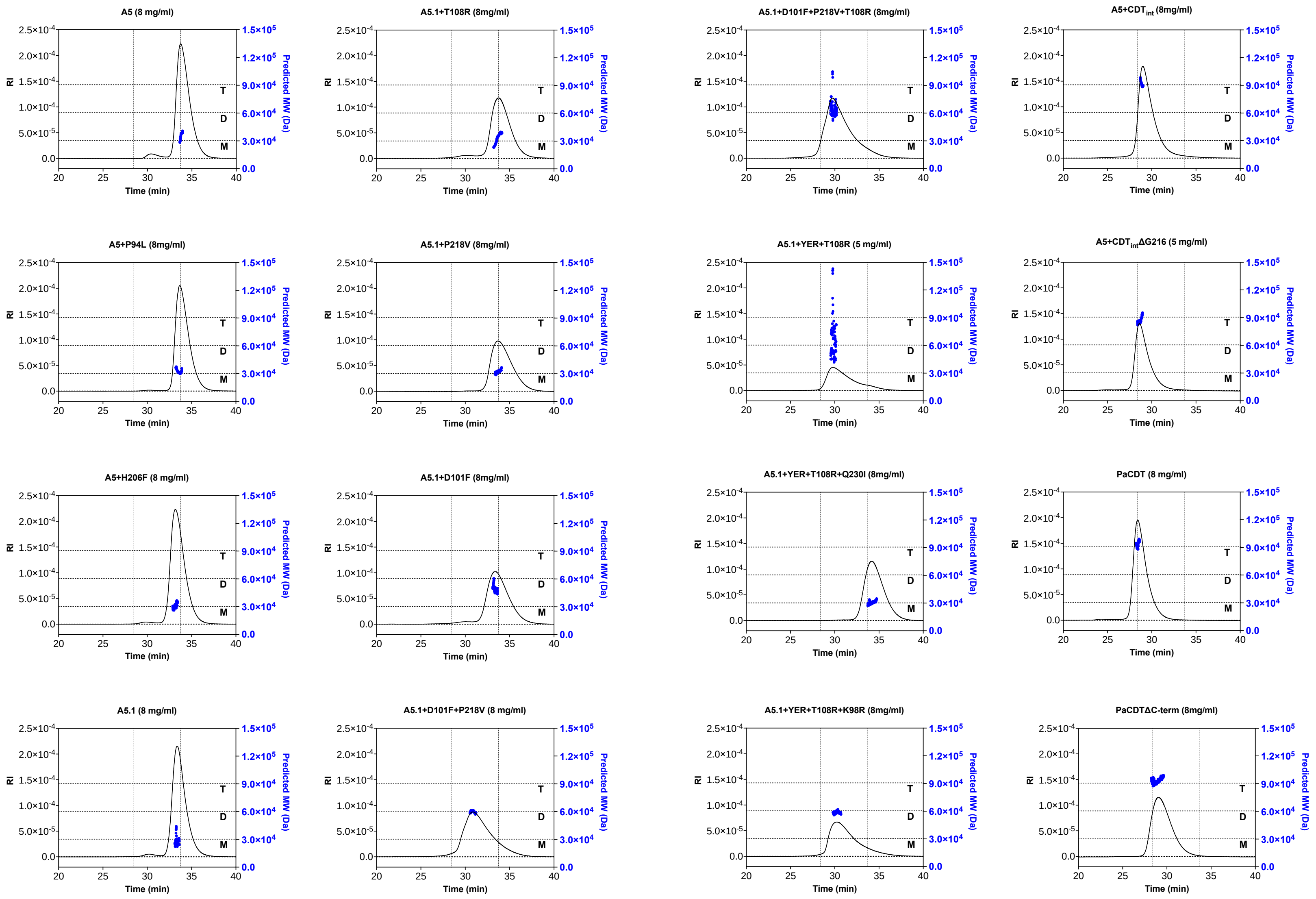


**Supplementary Figure 1. SEC-MALLS of the variants in this study at high concentrations (5-8 mg/mL).** Data from SEC-MALLS runs of variants using a Superdex 200 Increase 10/300 GL column, running at 0.5 mL/min in SEC buffer. Concentrations are noted in each panel. The black line represents the refractive index (RI), while the blue points represent the approximate molecular mass determined from light scattering. Horizontal lines corresponding to the expected molecular weights of the monomer (M), dimer (D), and trimer (T) are shown. Vertical lines corresponding to the elution volumes of AncCDT-5 and PaCDT are also shown for reference. Data is from a single run for each variant (n=1).

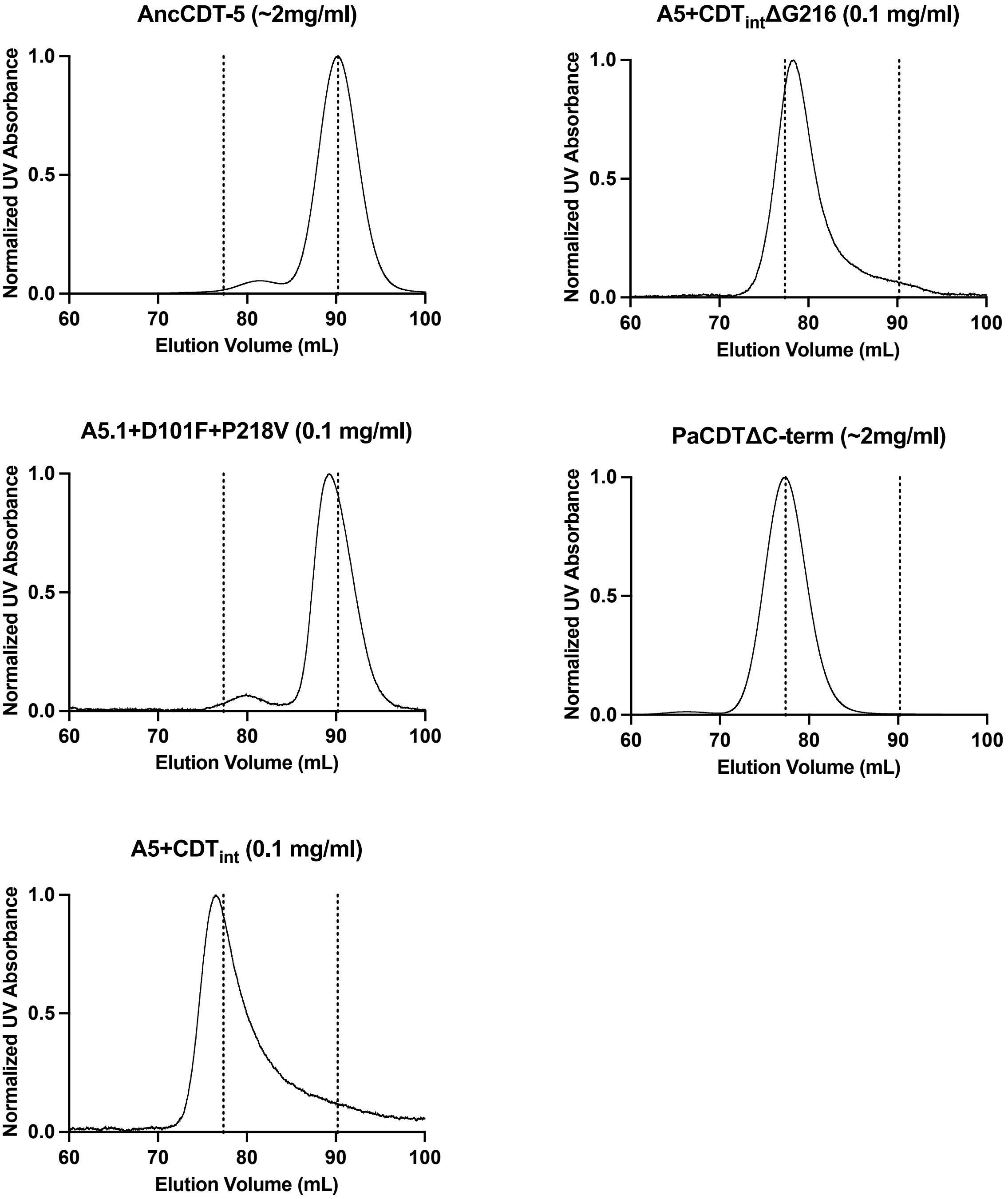


**Supplementary Figure 2. Normalized SEC chromatograms of A5.1+D101F+P218V A5+CDT$_{interface}$ and A5+CDT$_{interface}$ΔG216 at lower concentrations.** 4 mL of each sample was injected onto a HiLoad 16/600 Superdex 200 column (GE Healthcare) and eluted in SEC buffer. Vertical lines represent the elution volumes of PaCDTΔC and AncCDT-5.

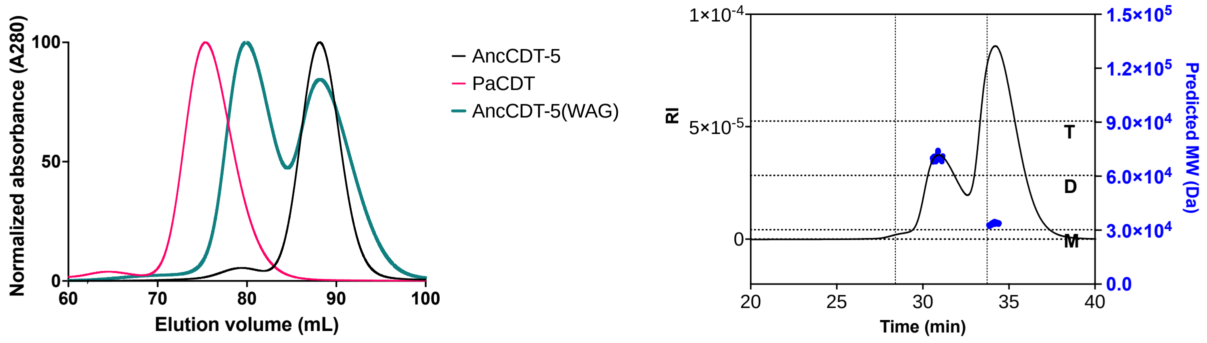


**Supplementary Figure 3. An alternate reconstruction of AncCDT-5, AncCDT-5(WAG), elutes with two peaks. (Left)** Overlay of representative chromatography traces from size-exclusion chromatography runs with PaCDT (orange), AncCDT-5 and AncCDT-5(WAG) on a HiLoad 16/600 Superdex 200 column. Plotted as normalized protein absorbance at 280 nm (based on the highest peak in each chromatogram). **(Right)** A SEC-MALLS run of AncCDT-5(WAG) (using the Superdex 200 10/300 GL size-exclusion column (GE Healthcare) in SEC buffer), with light scattering indicating that the larger peak likely corresponds to a dimeric form. A 100 μL sample was injected onto the column at approximately 8 mg/mL.

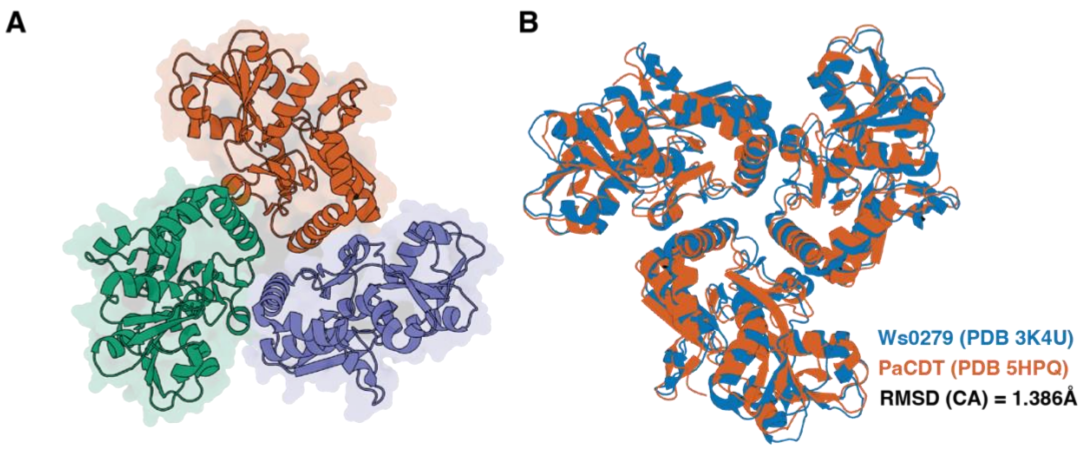


C

| | Chain 1 | | | | | | | | | | | | | Chain 2 | | | | | | | | | | |
|---|---|---|---|---|---|---|---|---|---|---|---|---|---|---|---|---|---|---|---|---|---|---|---|---|
| AncCDT-5 numbering (6WUP) | 92 | 94 | 97 | 98 | 101 | 108 | 206 | 208 | 215 | 216 | 218 | 221 | 222 | 16 | 17 | 20 | 218 | 219 | 222 | 226 | 229 | 230 | 233 | 234 |
| CDT numbering (6BQE) | 92 | 94 | 97 | 98 | 101 | 108 | 206 | 208 | 215 | - | 217* | 220 | 221 | 16 | 17 | 20 | 217 | 218 | 221 | 225 | 228 | 229 | 232 | 233 |
| PaCDT | I | L | Q | R | Y | R | F | E | R | - | E | K | R | L | D | L | E | A | R | Q | H | I | Q | S |
| Ws0279 | I | Q | N | L | N | V | Y | P | K | G | P | L | N | L | N | I | P | D | N | H | A | Q | H | D |
| 3K4U numbering | 72 | 74 | 77 | 78 | 81 | 88 | 186 | 188 | 195 | 196 | 198 | 201 | 202 | -4 | -3 | 0 | 198 | 199 | 202 | 206 | 209 | 210 | 213 | 214 |

D

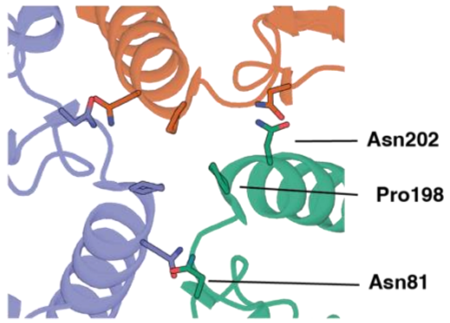


**Supplementary Figure 4. Ws0279 is a trimer in both solution and crystals. (A)** Ws0279 elutes from the HiLoad 26/600 Superdex 200 column as a trimer (elution volume of ~ 194 mL, calculated MW ~ 75 kDa, theoretical MW for dimer/trimer = 56 kDa/84 kDa; data not shown) and is a trimer in a crystal structure (PDB 3K4U). Ws0279 forms a similar trimeric arrangement as PaCDT in the crystal structure, with interactions primarily mediated by residues on the large sub-domains of the three chains, which are arranged around the point of three-fold symmetry. **(B)** Comparison of PaCDT (PDB 5HPQ) and Ws0279 (PDB 3K4U). **(C)** Comparison of residues at the interface positions in PaCDT and Ws0279. Apex positions are highlighted. **(D)** Residues at the apex positions in Ws0279 (PDB 3K4U) are shown as sticks.

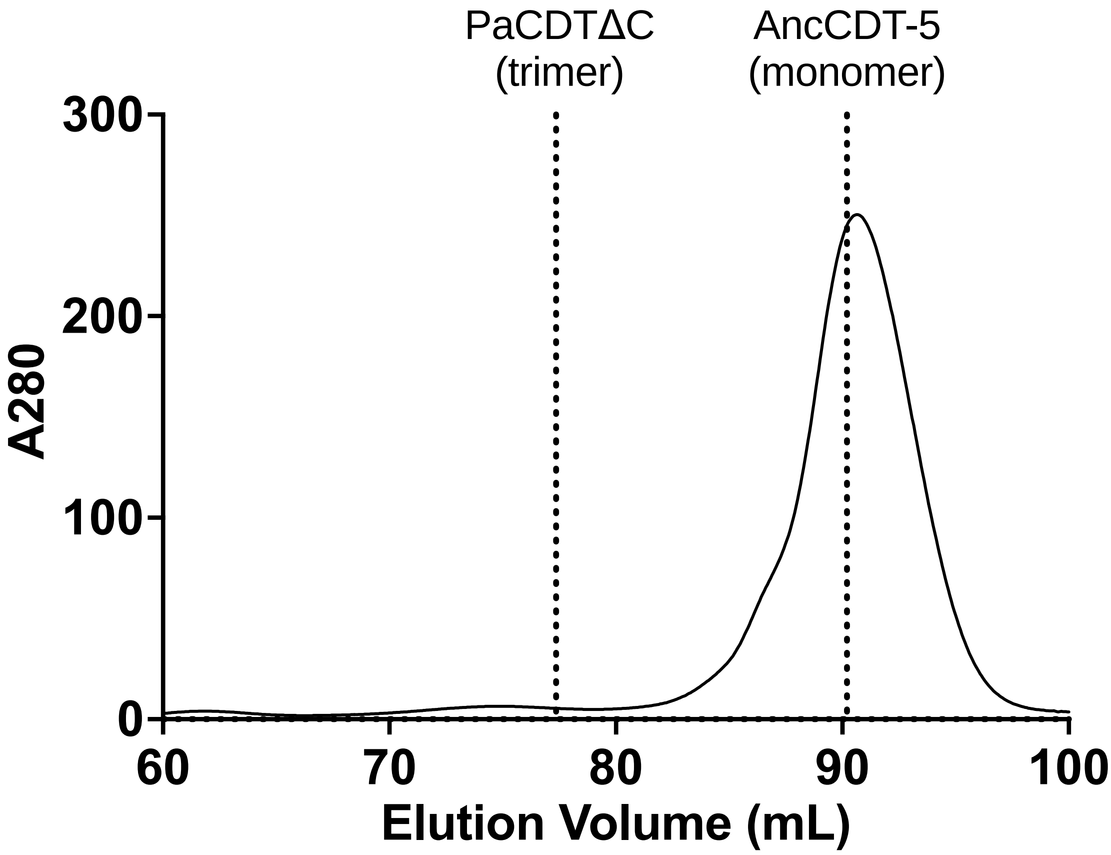


**Supplementary Figure 5. Ea1174 elutes as a monomer from SEC.** IMAC-purified Ea1174 was injected onto a HiLoad 16/600 Superdex 200 column (GE Healthcare) and eluted in SEC buffer. Vertical lines represent the elution volumes of PaCDTΔC and AncCDT-5.

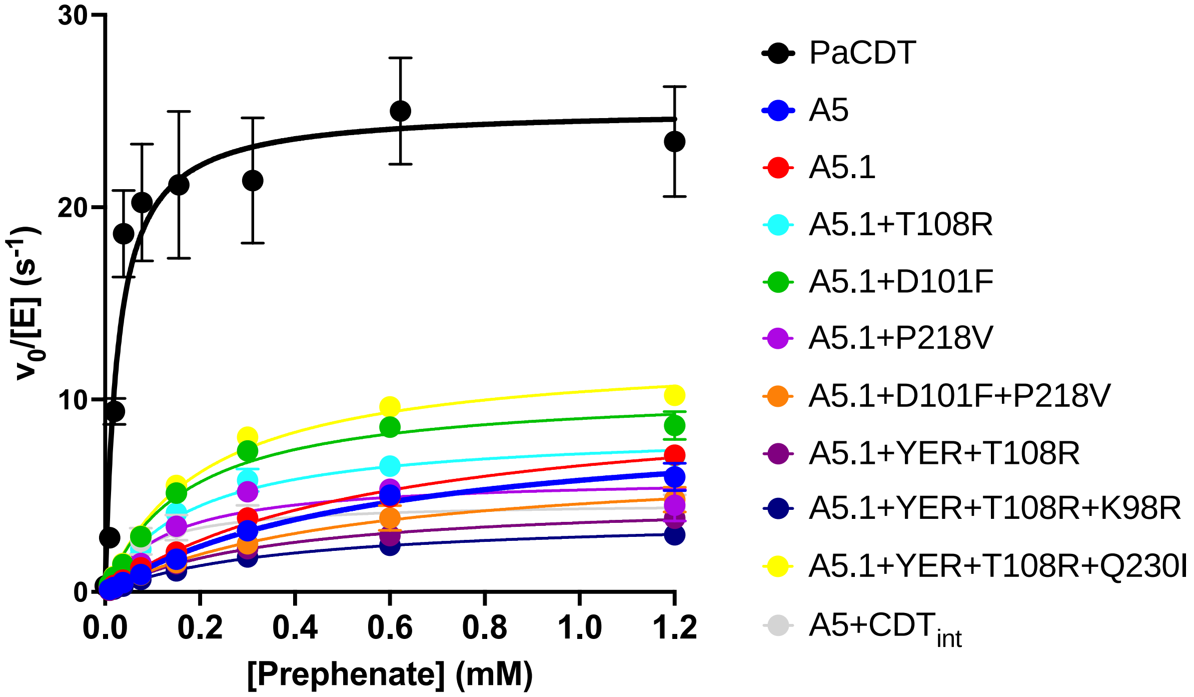


**Supplementary Figure 6.** Michaelis-Menton plots showing prephenate dehydratase activity of PaCDT, AncCDT-5 and key AncCDT-5 trimer-interface variants. Data represents the mean ± SEM (n ≥ 3 for all except A5.1+YER+T108R+K98R, for which n=2). Kinetic parameters are summarized in **Table I**.

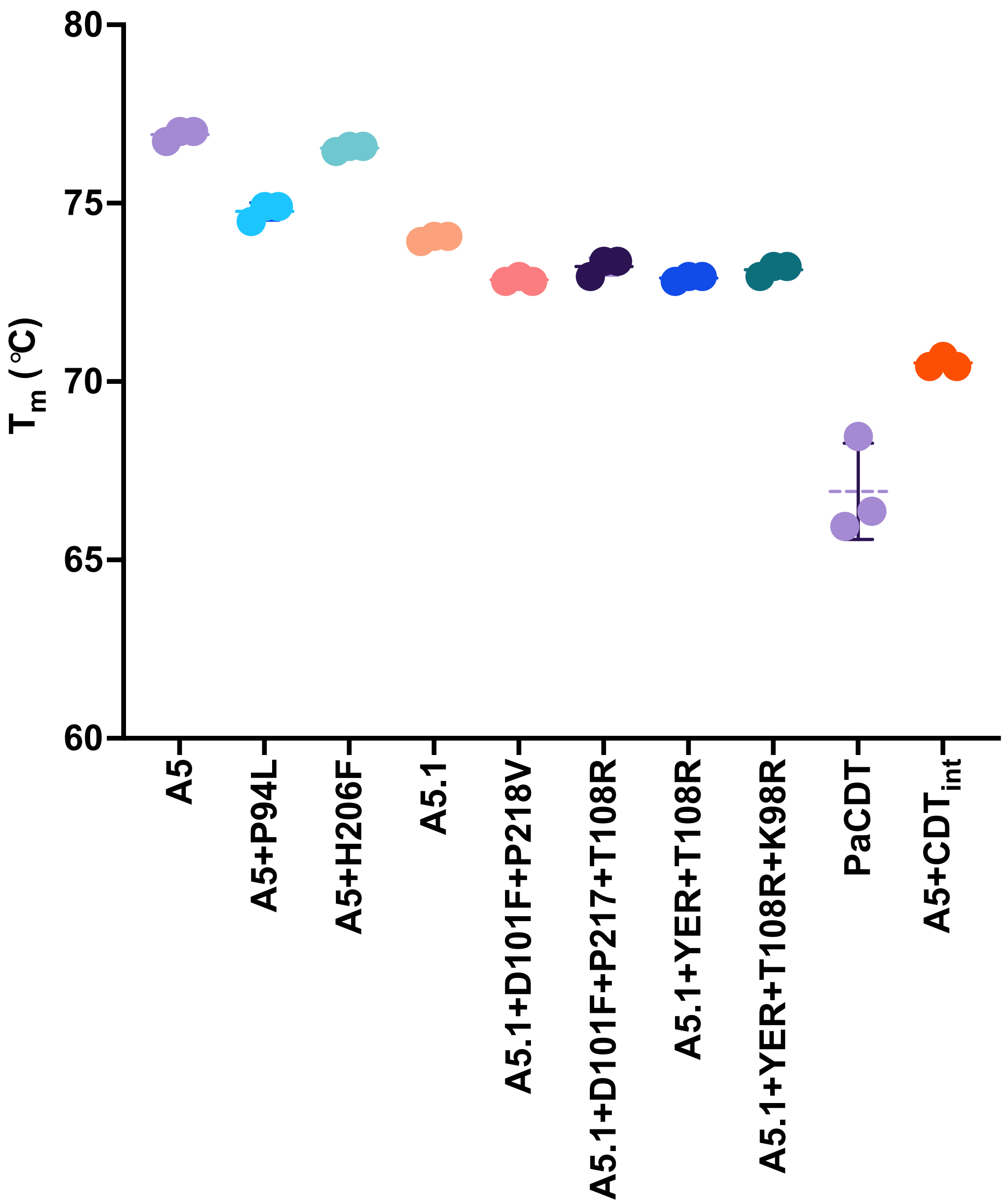


**Supplementary Figure 7. Thermostability of purified, His-tagged AncCDT-5, PaCDT and several AncCDT-5 trimer-interface variants.** Melting temperatures ($T_m$s) were determined based on the derivative of differential scanning fluorimetry traces. Each protein was tested as technical triplicates (n=3). The mean $T_m$ for each variant is shown as a horizontal line. Data also shown in **Supplementary Table II**.

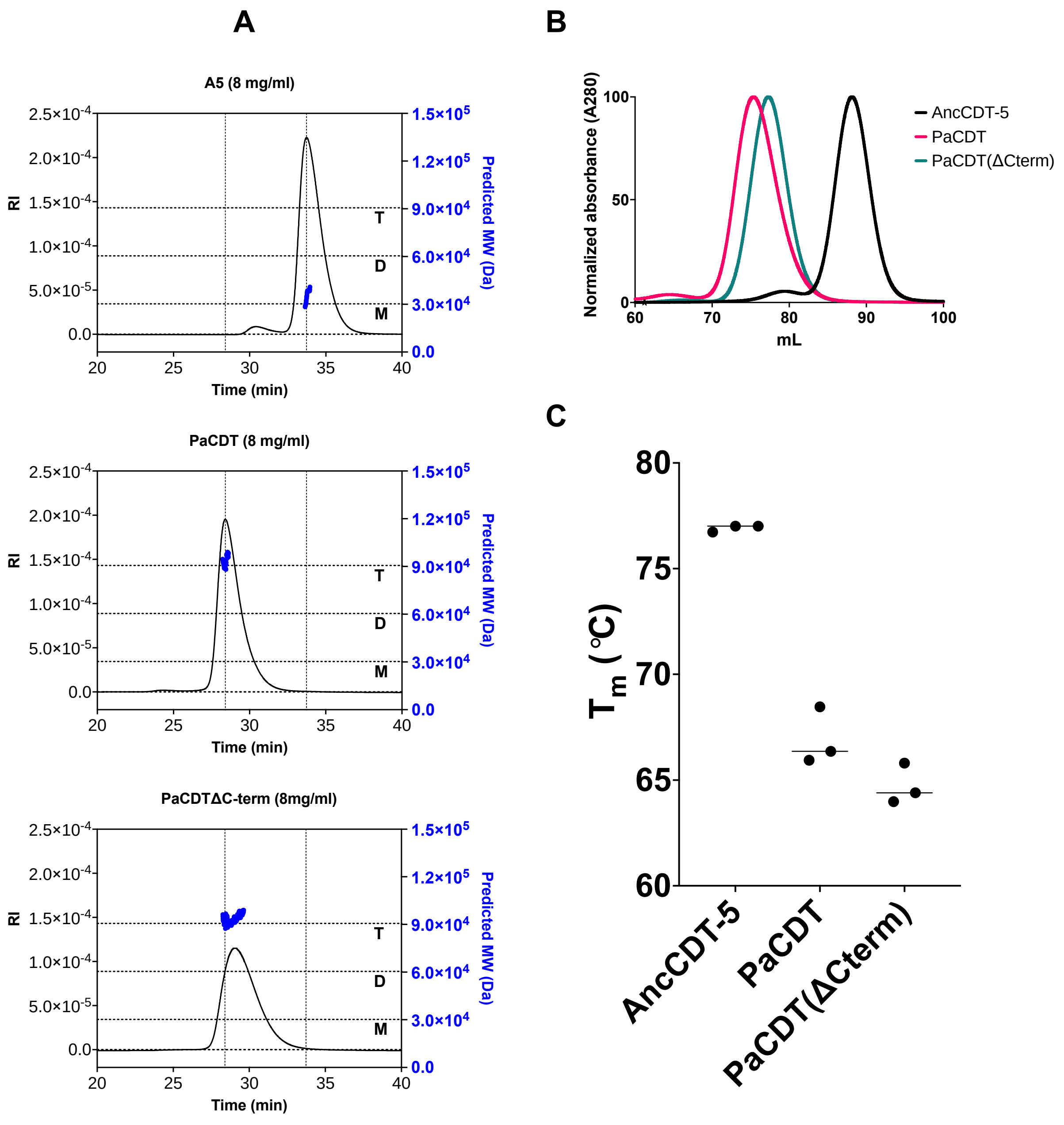


**Supplementary Figure 8. Truncation of the C-terminal extension from PaCDT.** **(A)** Representative SEC-MALLS runs for AncCDT-5, PaCDT and PaCDTΔC (using the Superdex 200 10/300 GL size-exclusion column (GE Healthcare) in SEC buffer). **(B)** Representative size exclusion chromatograms (normalized) for His-tagged AncCDT-5, PaCDT and the C-terminal-truncated PaCDT (PaCDTΔC) run on a HiLoad 16/600 Superdex 200 column. **(C)** Melting temperatures ($T_m$) for these variants (three technical replicates, n=3), as determined by differential scanning fluorimetry. The mean is represented by the horizontal line.

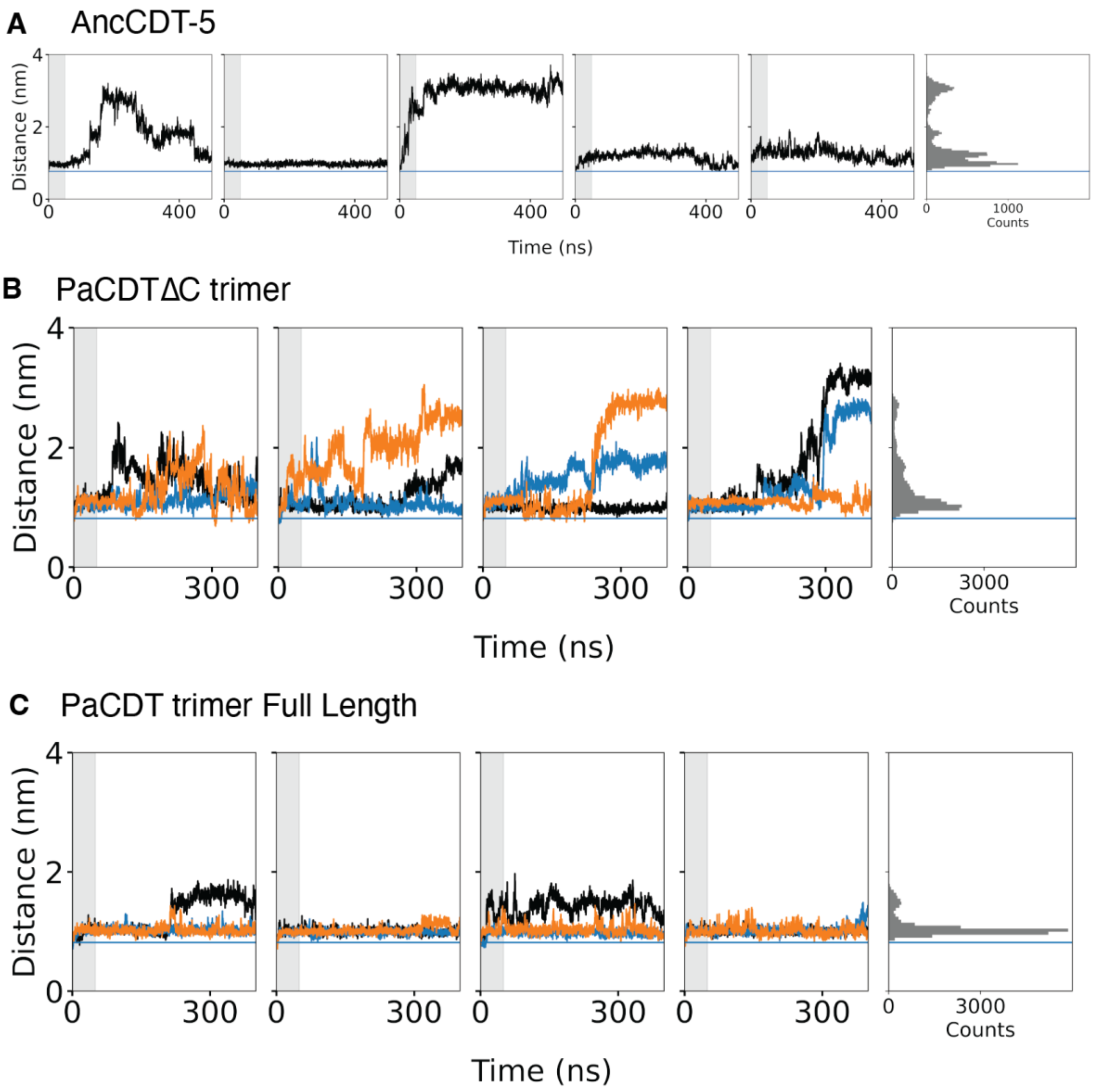


**Supplementary Figure 9. Molecular dynamics simulations of AncCDT-5, PaCDTΔC and PaCDT.** Plots showing how the distance between the alpha-carbon of Glu184 and Tyr33 varies during MD simulations. The blue horizontal line represents the Glu184–Tyr33 distance in the crystal structure of PaCDT (PDB 6BQE). Histograms on the right show the distribution of distances during the replicate simulations (the first 50ns of each simulation was omitted from histogram data). **(A)** Simulation initiated from chain A of the crystal structure of AncCDT-5 in which the C-terminal Lysine residue that was missing in the crystal structure was modelled (6WUP, residues 12 – 245 + Lys246). **(B)** Simulation initiated from trimeric structure of PaCDT in which the C-terminal extension had been removed. Individual chains represented by different colors. **(C)** Simulation initiated from trimeric structure of PaCDT.

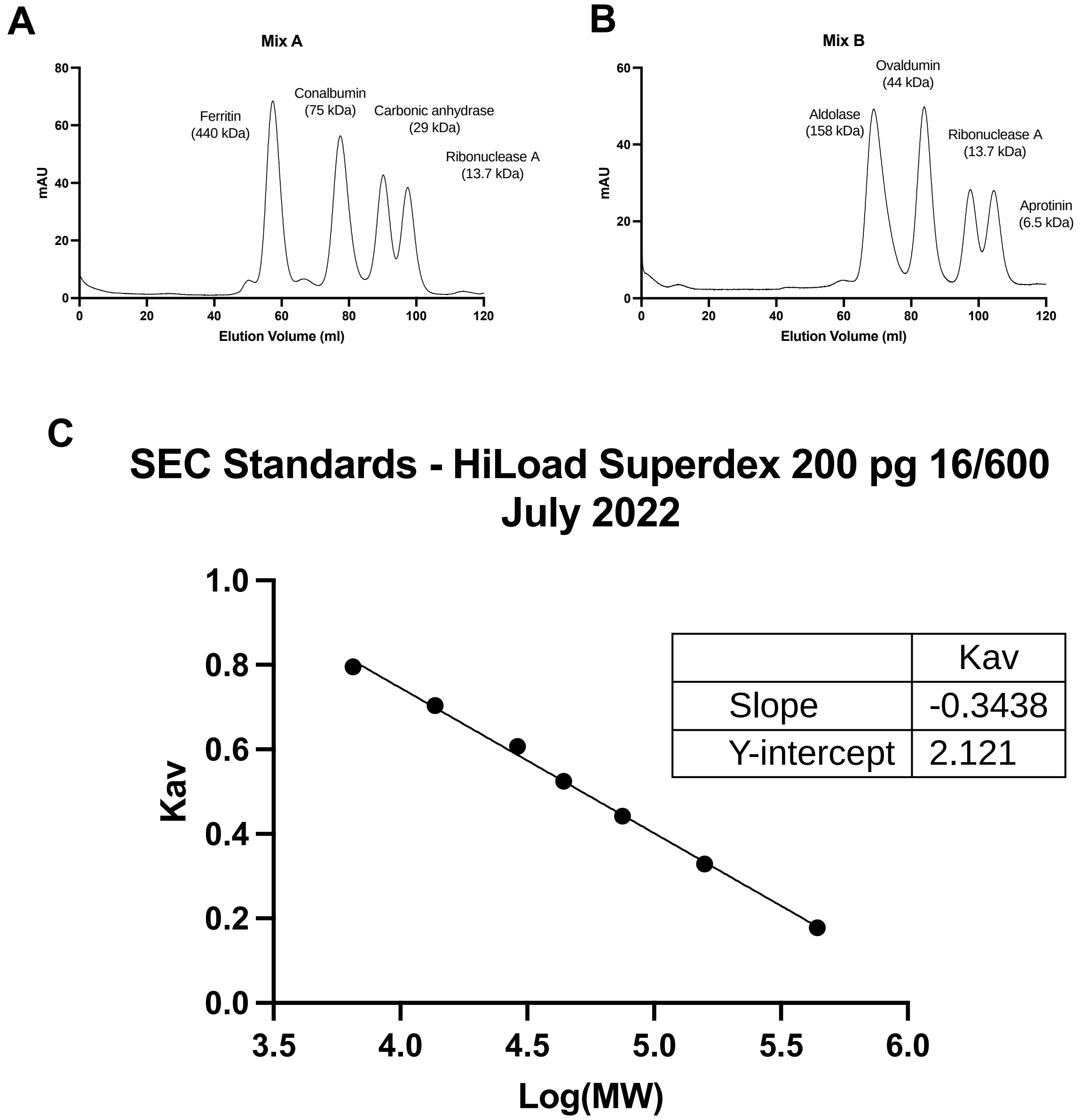


**Supplementary Figure 10.** Calibration of the HiLoad 16/600 Superdex 200 column (Cytiva) used for SEC-MALLS experiments using the Gel Filtration Calibration Kits (LMW, HMW) from Cytiva. Protein standards were injected on the column in two runs (**A, B**) to ensure peaks did not overlap. Column volume ($v_c$) was 120 mL and void volume ($v_0$) was determined to be 44 mL using Blue Dextran (not shown). Peaks were used for calculation of elution volume ($v_e$) and $k_{av}$ was according to $k_{av} = (v_e – v_0) / (v_c – v_0)$. All samples were run at 0.5 mL/min in SEC buffer. **(C)** Calibration curve that was used to estimate molecular masses of AncCDT-5 variants.

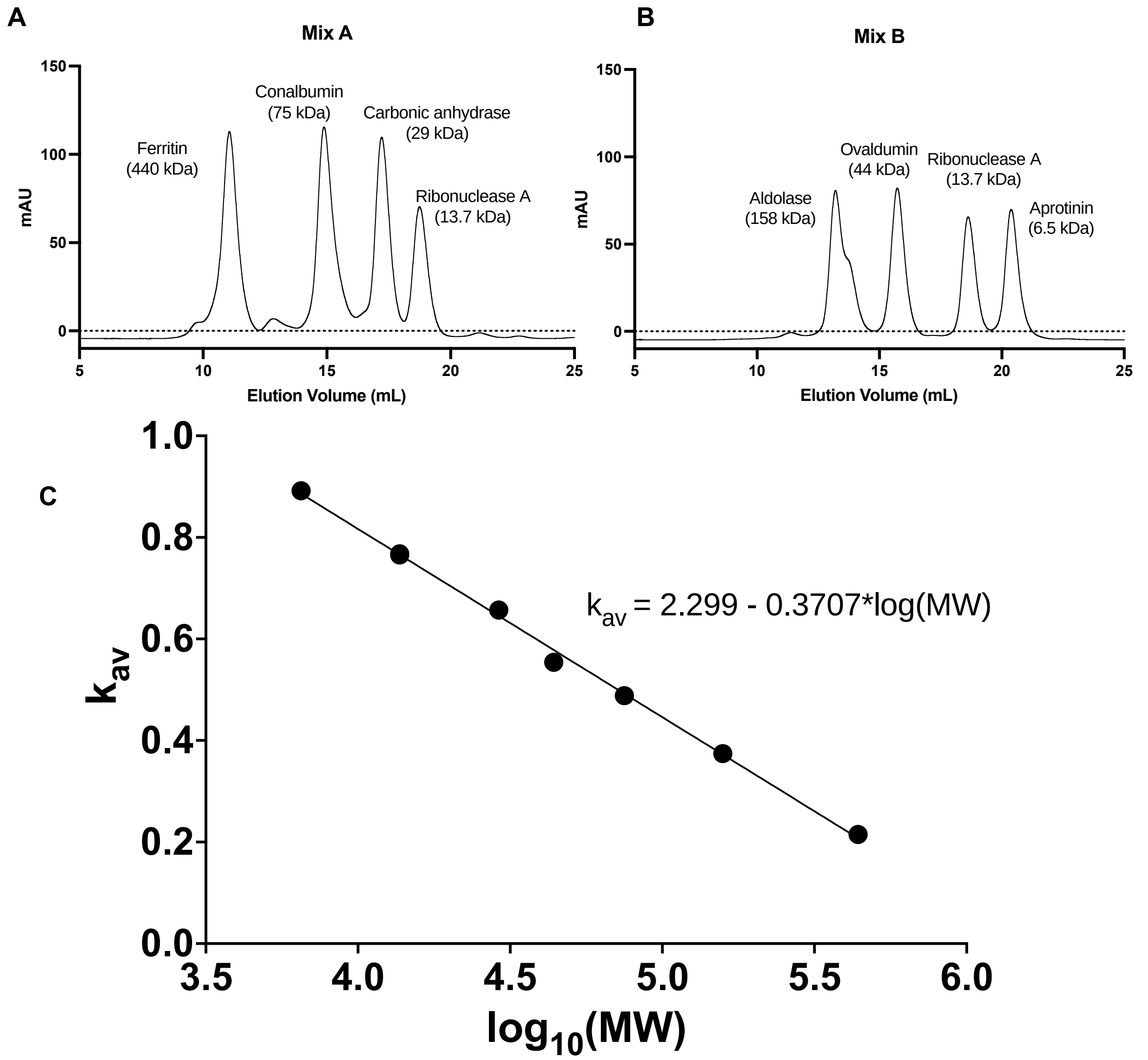


**Supplementary Figure 11.** Calibration of the Superdex 200 Increase 10/300 GL column (Cytiva) used for SEC-MALLS experiments using the Gel Filtration Calibration Kits (LMW, HMW) from Cytiva. Protein standards were injected on the column in two runs (**A, B**) to ensure peaks did not overlap. Column volume ($v_c$) was 22 mL and void volume ($v_0$) was determined to be 8.1 mL using Blue Dextran (not shown). Peaks were used for calculation of elution volume ($v_e$) and $k_{av}$ was according to $k_{av} = (v_e – v_0) / (v_c – v_0)$. Samples (100 µL) were run at 0.5 mL/min in SEC buffer. (**C**) Plot of $k_{av}$ vs $log_{10}$(MW) and linear line of best fit (equation shown on graph).

**Supplementary Table I. Interface residues positions identified from PaCDT crystal structures.** Residue positions identified in trimeric crystal structures (PDB IDs 6BQE, 3KBR, 5HPQ). Residues that are found in AncCDT-5 are shaded white, while those found in PaCDT are shaded dark grey. Those not found in either AncCDT-5 nor PaCDT are shaded light grey.

| | **Chain 1** | | | | | | | | | | | | | **Chain 2** | | | | | | | | | | |
|---|---|---|---|---|---|---|---|---|---|---|---|---|---|---|---|---|---|---|---|---|---|---|---|---|
| **AncCDT-5 res# (6WUP)** | **92** | **94** | **97** | **98** | **101** | **108** | **206** | **208** | **215** | **216** | **218** | **221** | **222** | **16** | **17** | **20** | **218** | **219** | **222** | **226** | **229** | **230** | **233** | **234** |
| **CDT res# (6BQE)** | **92** | **94** | **97** | **98** | **101** | **108** | **206** | **208** | **215** | **-** | **217*** | **220** | **221** | **16** | **17** | **20** | **217** | **218** | **221** | **225** | **228** | **229** | **232** | **233** |
| | | * | | * | * | * | * | | | | * | | * | | | | * | | * | | | * | | |
| **AncCDT-5 (A5)** | V | P | Q | K | D | T | H | E | R | G | P | K | A | L | D | M | P | A | A | Q | H | Q | Q | S |
| **AncCDT-5 (WAG)** | V | P | Q | K | D | R | H | E | R | G | P | K | A | L | D | M | P | A | A | Q | H | Q | Q | N |
| **Node A5a** | V | L | Q | K | F | T | F | E | R | G? | V | K | A | L | D | M | V | A | A | Q | H | Q | Q | S |
| **Node A5b** | V | L | Q | K | F | R | F | E | R | - | V | K | A | L | D | M | V | A | A | Q | H | Q | Q | S |
| **PaCDT** | I | L | Q | R | Y | R | F | E | R | - | E | K | R | L | D | L | E | A | R | Q | H | I | Q | S |
| **CDTchromob** | V | L | Q | K | L | N | F | Q | N | - | W | K | A | L | D | L | W | R | A | Q | S | Q | Q | D |
| **CDTralston** | V | L | Q | K | F | R | F | E | R | - | V | K | N | L | D | L | V | V | N | Q | R | Q | E | S |
| **CDTbradyrh** | I | L | Q | K | F | R | F | E | R | - | V | K | A | L | D | M | V | A | A | Q | H | M | E | N |
| **CDTmesorhi** | V | L | Q | K | F | R | F | E | R | - | M | K | A | L | D | V | M | A | A | Q | H | I | Q | D |

**Supplementary Table II. Thermostability of purified, His-tagged AncCDT-5, PaCDT and several AncCDT-5 trimer-interface variants.** Melting temperatures ($T_m$s) were determined based on the derivative of differential scanning fluorimetry traces. Each protein was tested as technical triplicates (n=3). This data is also shown graphically in **Supplementary Figure 7**.

| **Variant** | **Tm (°C)** |
|---|---|
| AncCDT-5 (A5) | 76.9 ± 0.2 |
| A5+P94L | 74.8 ± 0.2 |
| A5 + H206F | 76.5 ± 0.1 |
| A5.1 | 74.0 ± 0.1 |
| A5.1+D101F+P217V | 72.9 ± 0.1 |
| A5.1+D101F+P217+T108R | 73.2 ± 0.2 |
| A5.1+YER+T108R | 72.9 ± 0.1 |
| A5.1+YER+T108R+K98R | 73.1 ± 0.2 |
| A5+$CDT_{int}$ | 70.5 ± 0.2 |
| PaCDT | 66.9 ± 1.4 |

* Data is mean ± SEM of three technical replicates.

**Supplementary Table III. Details of each molecular dynamics system simulated and how they were prepared.**

| System Name | Initial PDB | Protein structure preparation & modelling | Ions in system ($Na^+/Cl^-$) | Production run length |
|---|---|---|---|---|
| AncCDT-5<br><br>6WUP_monomer | 6WUP | 6WUP was processed with PDB-REDO then loaded in Schrodinger Maestro. The terminal residue that was missing in the crystal structure (Lys246) was modelled using Schrodinger modelling tools followed by local minimization. Protein Preparation Wizard was used to remove small molecules (including acetate and waters). Missing residue atoms were filled. Added hydrogens. Optimized H-bonds. Restrained minimization performed. Protein termini were capped in gromacs. | 5 $Na^+$ | 500 ns x 5 |
| PaCDT_TRIMER<br>_TRUNCATED | 6BQE | 6BQE was processed with PDB-REDO. Removed small molecules (including acetate and waters). Residues 247-252 were deleted from each chain. Filled missing residue atoms. Added hydrogens. Optimized H-bonds. Restrained minimization performed. Protein termini were capped in gromacs. | 18 $Na^+$ | 500 ns x 4 |
| PaCDT_TRIMER<br>_CRYSTAL | 6BQE | 6BQE was processed with PDB-REDO. Removed small molecules (including acetate and waters). Filled missing residue atoms. Added hydrogens. Optimized H-bonds. Restrained minimization performed. Protein termini were capped in gromacs. | 15 $Na^+$ | 500 ns x 4 |

**Supplementary Table IV. Mutagenesis primers.** List of mutation primers used in this study to generate some of the AncCDT-5 trimer-interface variants.

| **Primer name** | **Primer Sequence** | **Parent Construct** |
|---|---|---|
| D101F_FW | AACCCTCGAACGTCAAAAAAAAGCCTTCTTTAGCGAGCCGTACATGACGGATGGC | A5.1 (pET28a) |
| D101F_RV | GCCATCCGTCATGTACGGCTCGCTAAAGAAGGCTTTTTTTTGACGTTCGAGGGTT | A5.1 (pET28a) |
| P217V_FW | ATTTGCTGCCGCGTGGTGATGTGGCGTTCAAAGCCTATGTGGACCAG | A5.1 (pET28a) |
| P217V_RV | CTGGTCCACATAGGCTTTGAACGCCACATCACCACGCGGCAGCAAATAAGC | A5.1 (pET28a) |
| T108R_FW | GCCGATTTTAGCGAGCCGTACATGCGTGATGGCAAAACGCCAATTGTCCGCTGC | A5.1 (pET28a) |
| T108R_RV | GCAGCGGACAATTGGCGTTTTGCCATCACGCATGTACGGCTCGCTAAAATCGG | A5.1 (pET28a) |
| Q229I_FW_GA | CGCTACGTTGATCAATGGCTGCACATTGCCATGCAATCAGGGACCTACCAG | A5.1+YER+T108R (pET28a) |
| Q229I_RV_GA | CTGGTAGGTCCCTGATTGCATGGCAATGTGCAGCCATTGATCAACGTAGCG | A5.1+YER+T108R (pET28a) |
| pET28_HalfGA_FW | CCAGCGCTTCGTTAATACAGATGTAGGTGTTCCACAGGGTAGC | any |
| pET28_HalfGA_RV | GCTACCCTGTGGAACACCTACATCTGTATTAACGAAGCGCTGG | any |
